# Cyclome: Large-scale replica-exchange dynamics of 930 cyclic peptide reveal thermal stability and critical metal-binding behavior

**DOI:** 10.64898/2026.04.08.717280

**Authors:** Karuna Anna Sajeevan, Hannah Gates, SR Vaishnavey, Curwen Pei Hong Tan, Riza Danurdoro, Julia Young, Ratul Chowdhury

## Abstract

Cyclic peptides are recognized as versatile scaffolds for therapeutic and functional applications due to their structural stability and resistance to degradation. Despite this promise, systematic analysis and prediction of their thermal stability remain limited by fragmented data resources, inadequate sequence comparison methods, and the lack of cyclicity-aware computational models. We provide a comprehensive, multi-scale computational framework to characterize cyclic peptides. First, we unified four fragmented public repositories of cyclic peptides into a single largest curated resource of 930 cyclic peptides, Cyclome930. This integrates cyclic topology, sequence, experimental structural coordinates, and source organism annotations into a consistently featurized dataset. Cyclome930 thus expands the dataset of annotated cyclic peptides by ∼3.4 fold (from 276 to 930). Second, we developed a novel cyclic sequence alignment algorithm that explicitly accounts for rotational symmetry and knot topology, enabling more accurate scoring of sequence similarity than conventional linear alignments. Third, we investigate the thermal stability of cyclic peptides using extensive all-atom replica-exchange molecular dynamics (100ns; REMD) simulations, allowing conformational sampling across 298 K – 400 K and track its stress tensors with increasing temperature. Finally, these simulation-derived thermo-stability metrics were used to train a machine learning model to predict cyclic peptide melting points from sequence and topology (STop2Melt). Crucially, the model introduces cyclicity-aware embeddings derived from ESMc representations coupled with cyclic offset vector, capturing the peptides’ knot topology. STop2Melt achieved strong predictive performance on held-out peptides and outperforms baseline methods that neglect cyclic structure. Finally, we scored Cyclome930 (cyclic ligands) for critical mineral metal binding using a multi-classifier model (CritiCL). To our knowledge, Cyclome930 represents the first effort in peptide literature to integrate physics-based temperature ramped simulations, cyclic sequence similarity scoring, machine learning for thermal stability prediction and scoring them for critical metal binding. Cyclicity-aware computational toolchains (<u>cyclome930.studio/</u>) provide a foundational resource for computational design of stable cyclic peptide prototype libraries thereby annotating and expanding genomic islands linked to critical mineral recovery.

## Introduction

Cyclic peptides are an emerging class of biomolecules that bridge the gap between small molecule drugs and larger protein therapeutics. They combine the desirable stability and membrane permeability of small molecules with the high specificity and potency of biologics.^1^ These closed-loop peptides, typically comprising a few to a few dozen amino acids, have garnered wide attention in drug development due to their unique pharmacological properties.^2^ Notably, cyclic peptides often exhibit potent bioactivities (e.g. analgesic,^3^ antibacterial^4^ and anticancer^2^), bind targets with high affinity and selectivity,^5,6^ and tend to have minimal off-target binding. Several favorable thermokinetic properties have made cyclic peptides invaluable in applications ranging from therapeutics and industrial *in vivo* biotechnology. For instance, cyclic peptide drugs are being explored to block “undruggable” protein–protein interactions in cancer and other diseases,^7^ leveraging their ability to precisely recognize and modulate biological targets *in vivo*.^8^ Indeed, the versatility and efficacy of cyclic peptides have positioned them as powerful tools in advancing modern pharmacology and biotechnology.^9^

Cyclic peptides are promising, hitherto untapped, thermostable, selective scaffolds for the selective recovery of critical minerals due to their structural stability, conformational rigidity, and diverse functional side chains capable of coordinating metal ions. Unlike linear peptides, cyclic peptides possess constrained backbones that enhance binding specificity,^6^ resistance to proteolysis^10^ (by exoproteases since some cyclic peptides do not offer a free terminus), and tolerance to wider thermo-chemical conditions,^11^ making them attractive candidates for use-inspired applications in metal ion recognition and separation. Many naturally occurring cyclic peptides and protein-derived motifs have evolved to coordinate metal ions through residues such as histidine, cysteine, aspartate, and glutamate, which can form stable coordination complexes with transition metals and rare-earth elements. While cysteine oxidation is often viewed as a liability in peptide systems,^12^ the conformational rigidity and tunable coordination environment of cyclic peptides make them particularly well suited for engineering highly selective chelators^13,14^ and can potentially be explored for critical minerals (CM) recovery. To this end, Cyclome930 provides a naïve prototypic rank-ordered library – with implications in clean energy technologies (Mn^2+^, Ni^2+^, Co^2+^, Ln^3+^ for electronic vehicles). Recent advances in computational protein design,^15,16^ peptide engineering,^17^ and machine learning have enabled the systematic exploration of peptide sequence space to identify sequences with enhanced critical metal-binding affinity and selectivity.^18^ Cyclic peptide scaffolds provide a versatile platform for tuning coordination geometry and binding pocket composition, allowing the development of biomolecular adsorbents with tunable kinetic lability, under aqueous conditions relevant to CM recovery processes. As the demand for sustainable and environmentally benign approaches for CM extraction grows, peptide and protein-based materials are increasingly being explored as biological pathways to CM recovery from dilute feedstocks. This offers potential advantages in selectivity and reduce environmental impact.^18–20^

The defining characteristic of cyclic peptides is the covalently closed backbone that results in a cyclized structure. This cyclization can occur through different chemistries and at various loci on the molecule. Common cyclization strategies include head-to-tail lactamization (forming a peptide bond between the termini), side-chain to tail linkage (an amide bond between an N-terminal amine and an internal side-chain carboxylate), head-to-side-chain linkage, or side-chain to side-chain linkages (disulfide bridges between cysteines).^21^ Accordingly, cyclic peptide knot topology in Cyclome930 is broadly dictated by the type of inter-residue linkage(s) along the peptide (**Figure 1**). They comprise five topological groups/ knot geometries: (1) end-to-end (e2e): cyclization occurs via a peptide bond between the N- and C-termini of the peptide chain, (2) side-to-end (s2e): involves a peptide bond between a side-chain functional group and either the backbone or side chain of a terminal amino acid, (3) side-to-side (s2s): formed through covalent linkages between side-chain functional groups, such as disulfide or ester bonds, (4) end-to-end with side-to-side (e2e+s2s): combines cyclization between the N- and C-termini with additional linkages between side-chain functional groups, and (5) Side-to-end with side-to-side (s2e+s2s): involves both a covalent bond between a side chain of a non-terminal amino acid and a terminal residue, and additional linkages between side-chain functional groups.

**Figure 1.**
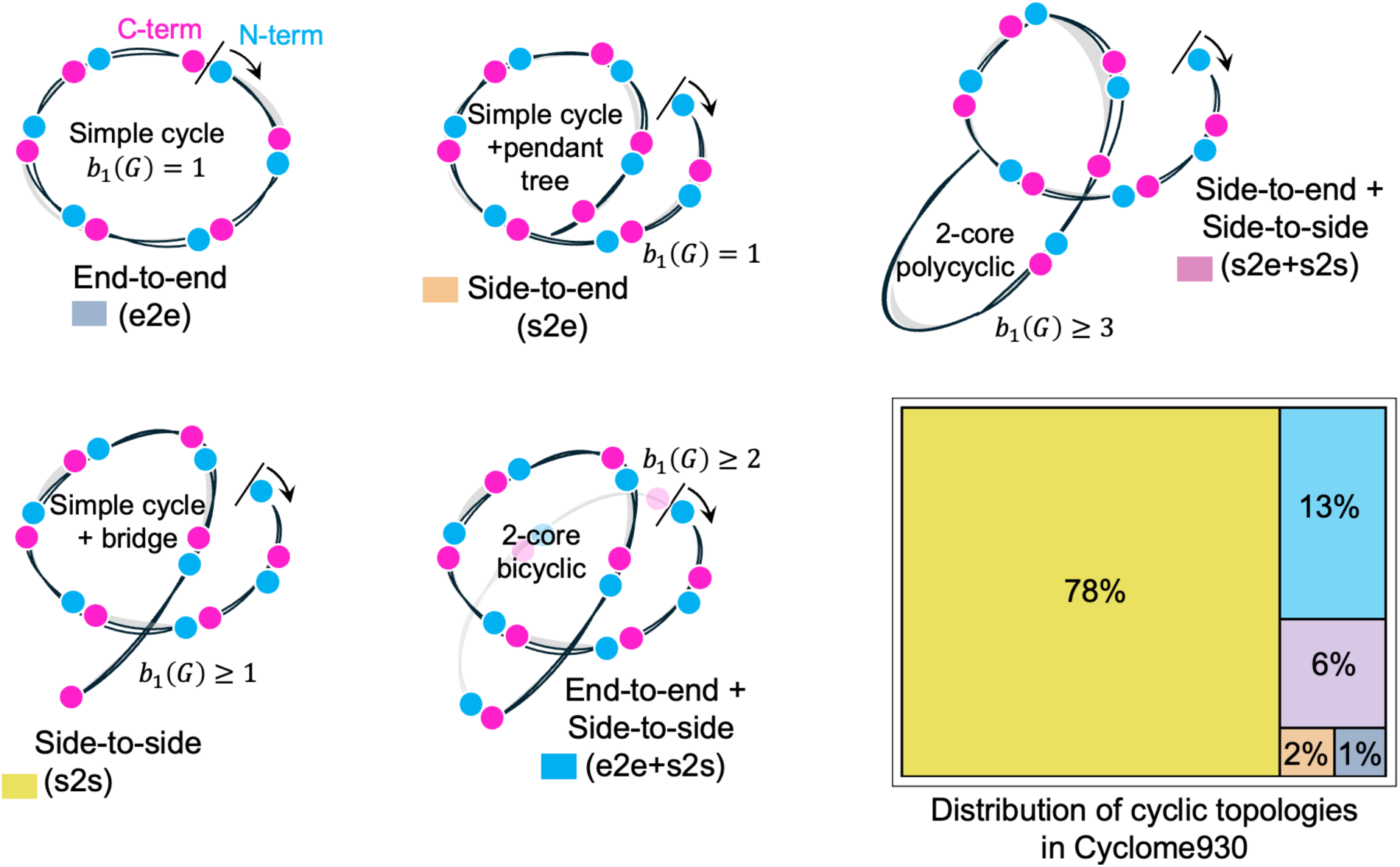
Topology classes of spatial graphs used in this work, indexed by first Betti number *b*_1_(*G*), which counts the number of independent cycles in each connected graph: (top left; **e2e (single macrocycle))**: A single simple cycle *C_n_* (*n* ≥ 11) with two designated termini, forming a 2-regular connected graph with first Betti number *b*_1_(*G*) = 1 and no bridges. All vertices have degree 2 and the unique cycle constitutes the entire 2-core, (top middle) **s2e (single loop with side junction))**: A unicyclic graph with *b*_1_(*G*) = 1 consisting of one simple cycle and a single pendant tree attached at exactly one cycle vertex. Exactly one cycle vertex has degree ≥ 3, all other cycle vertices have degree 2, and the edges off the cycle form a single tree terminating in an end vertex. (bottom left; **s2s (loop with internal bridge))**: A unicyclic graph with *b*_1_(*G*) = 1 that contains at least one bridge whose endpoints both have degree ≥ 2. The unique cycle remains intact upon removal of this bridge, while a nontrivial tree-like substructure is detached, yielding an internal “bridge” within the loop rather than a second independent cycle. (bottom middle; **e2e + s2s (bicyclic spatial graph))**: A connected graph with cycle rank *b*_1_(*G*) = 2, representing two independent macrocycles in the same component. The 2core is bicyclic, comprising-either a single block with two linearly independent cycles in *H*_1_(*G*, ℤ) or two cyclic blocks connected by bridges; any remaining edges are trees attached to this bicyclic backbone. (top right; **s2e + s2s (multi-cycle spatial graph)**: A connected, polycyclic graph with *b*_1_(*G*) ≥ 3. The 2-core is neither unicyclic nor merely bicyclic but instead contains three or more independent cycles and/or multiple cyclic blocks linked by bridges, with additional tree-like branches attached to this multicycle core. In addition, a hierarchical chart shows the % of each type of topology in Cyclome930.

Classifying cyclicity by knot topology **(Figure 1)** provides a rigorous way to connect sequence–structure space to thermodynamic stability. In particular, the first Betti number *b*_1_(*G*)of the spatial graph associated with a cyclic polypeptide backbone counts the number of independent cycles, thereby distinguishing simple macrocycles ( *b*_1_= 1 ), bicyclic motifs ( *b*_1_ = 2 ), and higher-order polycyclic knotted states ( *b*_1_ ≥ 3 ). By organizing structures according to *b*_1_(*G*) and the arrangement of bridges and side-junctions, Cyclome930 defines a topological state space in which changes in melting temperature can be interpreted as responses to how free energy is distributed across distinct cyclic subgraphs.

Because melting temperature reflects the balance between enthalpic stabilization and entropic costs, understanding how each topological class stores and redistributes free energy is critical. In tightly knotted or multi-cyclic backbones, increased *b*_1_(*G*) generally implies more constraints on chain configurational entropy and more pathways for distributing mechanical and electronic strain under heating. Thus, when local total free energy increases (through thermal input), the way knots tighten, slip, or delocalize stress depends systematically on the underlying cycle structure defined by Betti number and cycle connectivity. Cyclome930 therefore offers a fundamentally grounded theoretical explanation of melting temperature: it links macroscopic thermal transitions to the homological features of the peptide’s spatial graph.

The scope of this paper is to assemble the ingredients needed to open several new lenses on cyclic polypeptide science. In this work, we lay the foundations to connect how thermal stress tensors evolve over time across the cyclic backbones. Future endeavors could leverage these fields to calculate critical mineral binding across 298 − 400 K. Because each metal-binding configuration is indexed by its knot class and *b*_1_(*G*), the resulting dataset can be reused to learn how critical mineral–metal binding changes with temperature. Practically, this involves applying thermal corrections to the effective spring constants assigned to every bond in the system including inter–amino acid peptide bonds and the bridge bonds that define intra- and inter-cycle connectivity. By grouping these springs according to the topological types introduced in Cyclome930 (e.g., e2e, s2e, s2s, and their combinations), we obtain a parametric map from Betti-number–resolved graph topology to temperature-dependent mechanical response, metal affinity, and ultimately melting behavior (**Figure 2**).

**Figure 2.**
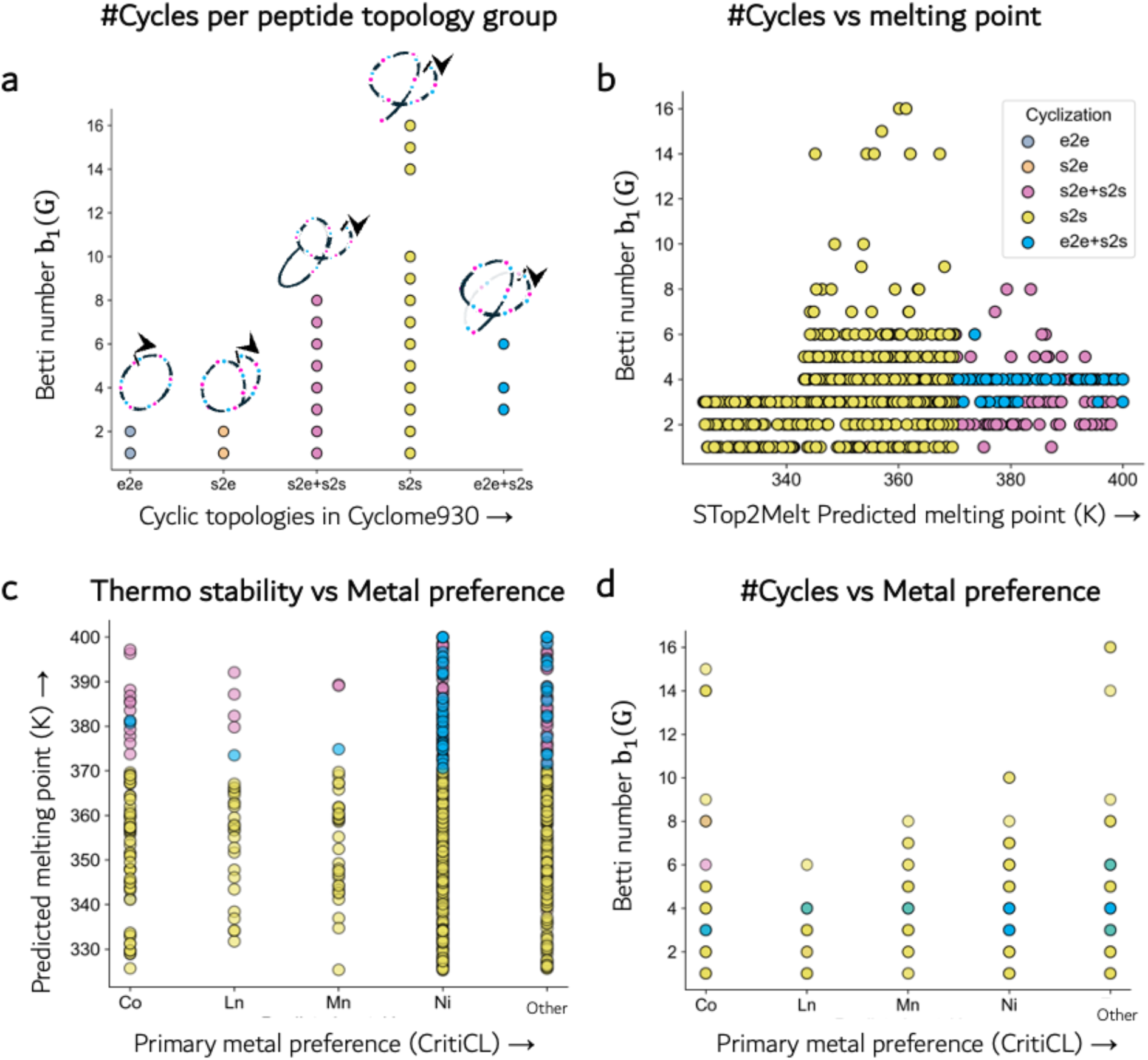
Topological complexity and melting behavior of cyclic peptides across cyclization classes and predicted metal-binding categories. **(a)** Distribution of the first Betti number *b*_1_(*G*), representing the number of independent cycles in the residue contact graph, across different cyclization classes (e2e, s2e, s2e+s2s, s2s, e2e+s2s). Schematic cartoons illustrate representative topological motifs for each cyclization type. Increased structural constraints introduced by additional crosslinks generally correspond to higher first Betti numbers. **(b)** Relationship between predicted melting temperature (STop2Melt) and topological complexity *b*_1_(*G*). Points are colored by cyclization class, highlighting that peptides with side-to-side disulfide connectivity (s2s) tend to populate higher first Betti numbers and span a broad thermal stability range. **(c)** Distribution of predicted melting temperatures for cyclic peptides grouped by predicted metal-binding class (Co^2+^, Ln^3+^, Mn^2+^, Ni^2+^, and other metals). Distinct cyclization classes show varying stability ranges, with highly constrained architectures generally exhibiting higher predicted thermal stability. **(d)** Variation of topological complexity *b*_1_(*G*) across predicted metal-binding classes, indicating that peptides predicted to bind different metals occupy distinct regions of the topology–stability landscape. Colors correspond to cyclization classes as defined in panel (a).

A major advantage of cyclization is the enhancement of peptide stability. Lacking free N- and C-termini, cyclic peptides are inherently more resistant to exopeptidases and other degrading enzyme.^22^ The rigid ring structure pre-organizes the peptide, reducing the entropy cost upon binding and often locking the molecule into a stable conformation.^23^ Consequently, cyclic peptides tend to be significantly more stable than similarly sized linear peptides under physiological/ harsh physical or chemical stress. For example, cyclotides are notoriously resistant to degradation – the prototypical cyclotide kalata B1 remains folded and bioactive even after boiling and exposure to denaturants.^24^ This exceptional stability is attributed to the cyclic cystine-knot motif, which imparts rigidity against thermal, chemical, and proteolytic challenges.^25^ Likewise, lasso peptides are generally heat-stable up to a threshold; heating beyond that can irreversibly unthread the lasso into a less stable cyclic form.^26^ Such robustness is highly desirable in drug development, as it can translate to improved shelf-life, oral availability, and efficacy *in vivo*. Thermal stability is an important quality for peptide therapeutics: a drug that remains folded at elevated temperatures is less likely to aggregate or lose function during manufacturing and storage. It can also indicate a high energetic barrier to unfolding, often correlating with stability in the presence of denaturants or proteases. Despite its importance, there is a dearth of systematic data on the thermal stability of cyclic peptides in the literature – thus there is no established clear topology-to-melting mapping for a given cyclic peptide sequence (hereafter, STop2Melt). Most studies have focused on individual cases (e.g. cyclotides or lasso peptides)^24,26^ and broad surveys or predictive models of cyclic peptide melting temperatures or structural stability are largely missing. Filling this knowledge gap is crucial, as understanding how sequence and structure influence thermal resilience could guide the design of new cyclic peptides with improved stability profiles.

Cyclic peptide information has historically been scattered across general protein/natural product databases and individual studies, hindering efficient featurization and analysis. For example, CyBase, a database created for cyclic peptides/proteins, is heavily focused on plant cyclotides and a few related families, with cyclotides comprising ∼75% of its entries.^27^ It also lacks systematic integration of structural information though some entries are linked to PDB structures. Overall, prior resources either covered only subsets of the cyclic peptide chemical space or were not maintained in a way to facilitate comprehensive analysis. Recently, the importance of a unified resource has been acknowledged: Liu *et al.* highlighted that existing public databases contained cyclic peptide data in a dispersed and non-standardized manner, weakening their utility for drug discovery.^28^In response, efforts like the CyclicPepedia knowledge base assembled over 8,700 cyclic peptides, both naturally occurring and synthetic, along with annotations on their structures, sources, and activities. Nevertheless, it has only 276 cyclic peptides with structural information. This underscores the community’s recognition that a comprehensive, well-curated cyclic peptide database is essential to drive further research and development. Building on this momentum, our work aims to significantly expand and enrich the available data, providing an even more complete resource that also incorporates extensive structural information.

Analyzing cyclic peptide sequences poses unique challenges because of their circular nature. Standard bioinformatics tools for sequence alignment and comparison assume linear sequences with defined start and end points; applied naively to a cyclic sequence, the choice of linear starting point is arbitrary and can lead to incorrect alignment conclusions. Two identical (or highly similar) cyclic peptides may appear dissimilar if one is “rotated” relative to the other in sequence form.^29^ Therefore, specialized alignment algorithms or sequence representations are needed to recognize equivalences under rotation. Grossi et. al has noted that incorporating cyclic permutation invariance is necessary for accurate comparison of circular sequences.^29^ In this study, we address this issue by developing a *cyclicity-aware sequence alignment* method. By conceptually permuting sequences and optimizing alignments over all rotations (or by using cyclic string-matching techniques), our approach can correctly align all five topologies of cyclic peptides independent of their linear starting positions. Likewise, for machine learning (ML) applications, we introduce a cyclicity-aware sequence embedding, which ensures that peptides with the same circular sequence receive the same representation regardless of how their sequence is rotated. This idea is inspired by recent advances in structure prediction for cyclic peptides, where positional encodings were modified to be rotation-invariant to successfully apply protein large language models like AlphaFold2.^30^ By embedding rotational invariance into our feature engineering, we enable ML models to learn from cyclic peptide data without being confounded by sequence indexing artifacts.

Here, we present an integrated computational framework to advance cyclic peptide research. First, we have compiled a comprehensive database of cyclic peptides, Cyclome930 aggregating sequences and structures from multiple sources and systematically curating their annotations. This database expands previous efforts from 276 unique cyclic peptides with structural information to 930, providing a foundation for large-scale analyses and model building. Second, we implement a novel cyclic sequence alignment algorithm that can identify sequence similarities among cyclic peptides more effectively than linear alignments. Third, we investigate the thermal stability of cyclic peptides through extensive replica-exchange molecular dynamics (REMD) simulations. REMD allows us to sample peptide conformations over a range of temperatures, from which we can infer folding/unfolding behavior and stability parameters for representative peptides. These simulation-derived stability metrics are then used to train and validate an ML model that predicts thermal stability for cyclic peptides *in silico*. Crucially, our ML model utilizes a cyclicity-aware embedding of sequences, to represent the peptides’ circular nature. Finally, we screened Cyclome930 for their binding to critical minerals to screen their potential for critical mineral capture. This work represents the first attempt to predict thermal stability of cyclic peptides by combining physics-based simulations with machine learning and screening them for critical mineral capture.

## Results

### Database composition

To establish a high-quality and comprehensive cyclic peptide database, we compiled all available cyclic peptide entries with sequence and/or structural annotation from RCSB PDB, ConoServer, CyBase, and CyclicPepedia.^27,28,31,32^ These resources contain 211,460 PDB entries, 4,832 ConoServer peptides, 1,936 CyBase peptides, and 7,739 CyclicPepedia peptides, respectively (**Table 1**). After harmonizing these repositories, we filtered specifically for entries with experimentally determined 3D structures, yielding 1,059 cyclic peptide structures (**Figure 3a**). Because several PDB entries contained more than one peptide chain, chain-level parsing expanded the dataset to 1,734 individual chains. Next, we removed redundancy at the sequence level to obtain unique cyclic peptide chains, while retaining the full mapping of all PDB structures corresponding to each sequence. This produced a curated set of 930 nonredundant cyclic proteins/peptides, hereafter termed Cyclome930.

**Figure 3.**
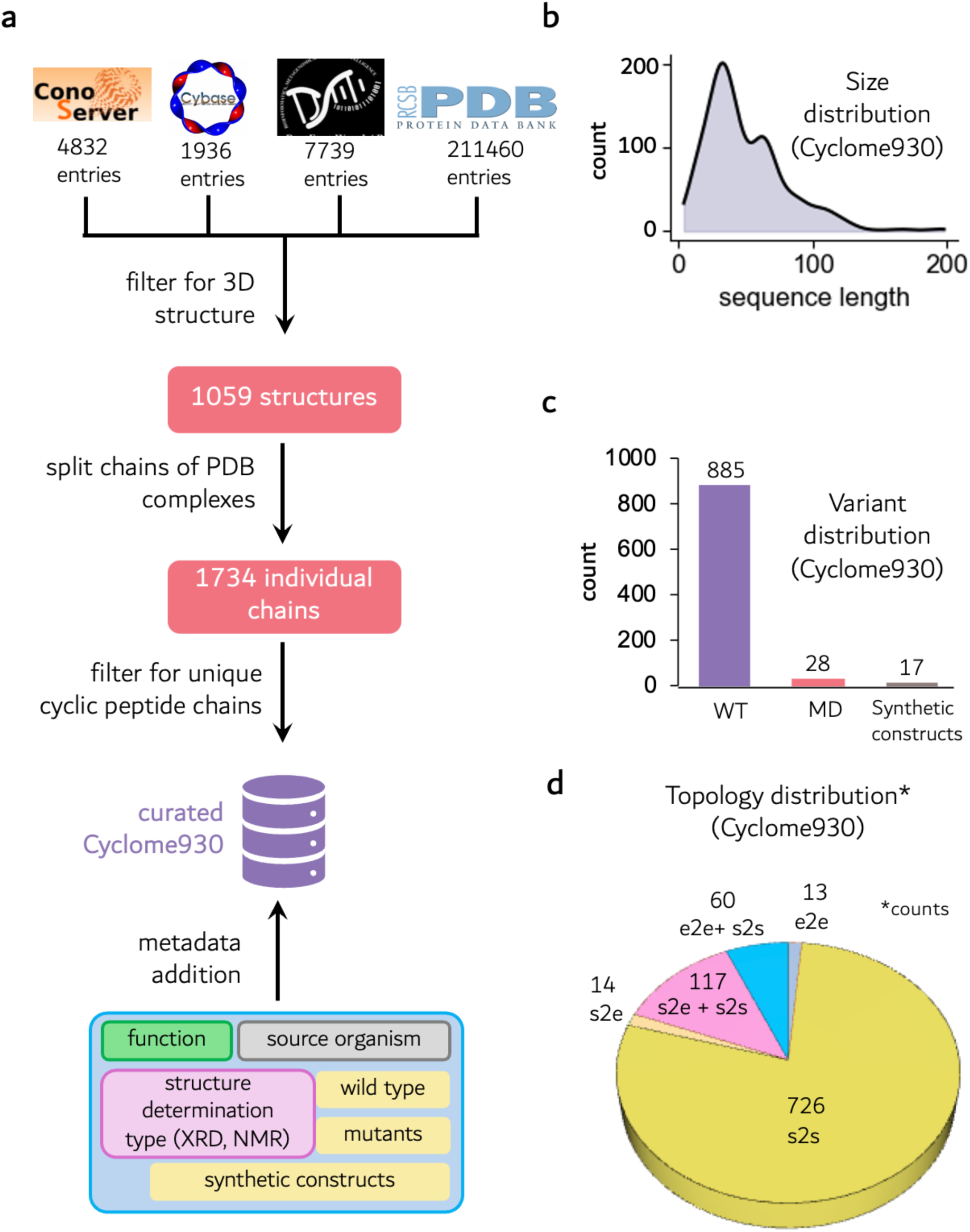
Construction and overview of the Cyclome930 cyclic peptide structural dataset. (**a)** Cyclic peptide structures were curated from multiple public databases including ConoServer,^31^ CyBase,^27^ CyclicPepedia^28^ and the RCSB Protein Data Bank (PDB).^32^ The initial collection comprised 1059 structures, corresponding to 1734 individual peptide chains after separating multichain entries. Following redundancy removal and sequence-based filtering, a curated dataset termed Cyclome930 was generated. (**b)** The distribution of peptide sequence lengths shows that most cyclic peptides fall within length 10-40. **(c, d)** Additional dataset statistics illustrate wild-type, mutant (MD), or synthetic constructs, Cyclization class representatives in Cyclome930. Together, these curated features provide a comprehensive structural resource for downstream computational analysis and machine learning studies of cyclic peptides.

**Table 1.** Comparison of Cyclome930 with existing cyclic peptide databases. Summary of sequence counts, experimentally determined structures, source organisms, and synthetic constructs across CyBase, ConoServer, CyclicPepedia, and Cyclome930. Cyclome930 provides the largest set of structurally validated cyclic peptides, with 930 unique chains, broad organismal coverage, and inclusion of 17 synthetic constructs.

|  | Cybase<sup>27</sup> | Conoserver<sup>31</sup> | CyclicPepedia<sup>28</sup> | Cyclome930 |
| --- | --- | --- | --- | --- |
| Sequence | 1936 | 4832 | 7739 | 930 |
| 3D-Structure | **83** | **247** | **276** | **930** |
| Source organism | 180 | 126 | 764 | 244 |
| Synthetic construct | - | 83 | 1434 | 17 |

Cyclome930 represents the largest cyclic peptide database to date with both sequence and structural annotation, exceeding prior datasets such as CyBase,^27^ ConoServer,^31^ and CyclicPepedia^28^ in the number of experimentally resolved 3D structures (**Table 1**). The database spans 244 source organisms and includes 17 synthetic cyclic constructs that were preserved as distinct sequence entries. Structure depositions range from 1980 to 2025, with 413 structures determined by X-ray diffraction and 556 by multidimensional NMR, reflecting broad methodological coverage. The curated dataset also allows metadata-level analyses (**Figure 3b–d**). The sequence length distribution is heavily skewed toward peptides of 10–40 residues. Cyclome930 includes 885 wild-type peptides, 28 mutant constructs and 17 synthetic peptides (**Figure 3c**). Classification by cyclization pattern showed that Cyclome930 has representatives from all five categories: (i) *e2e* peptides (13), which cyclizes solely via head-to-tail backbone ligation, (ii) *s2s* peptides (726), defined by one or more side-chain–to–side-chain covalent linkages such as disulfide bonds, (iii) *s2e* peptides (14), formed exclusively through side-chain of a nonterminal residue to-terminal residue cyclization, (iv) *s2e+s2s* (117), where side chain of a nonterminal residue links to terminal residue, along with other sidechain to side chain covalent linkages, (v) *e2e+s2s* (60) where end to end backbone ligation is combined with side chain to side chain linkage (**Figure 3d**). The **Cyclome930** database containing all sequences, structures, and metadata annotations is publicly available for download at: https://cyclome930.studio/. Cyclome930 studio is designed to support systematic sequence analysis, structural characterization, and machine-learning model development. The database serves as a foundational resource for investigating cyclic peptide properties, including stability and folding behavior, and for developing cyclicity-aware computational methods.

### Cyclicity-aware sequence similarity predictor

Determining sequence similarity for cyclic peptides requires special consideration of cyclization type, as conventional linear similarity calculators often overlook the cyclic nature of these molecules, leading to systematic underestimation of similarity scores. To address this limitation, we developed a cyclization-specific sequence similarity calculation framework. For end-to-end (**e2e**) cyclized peptides (**Figure 5a**), the template sequence is duplicated and concatenated in the N→C direction to generate a *true template* (T-template, **Figure 4**). Similarity between a query sequence and the T-template is then evaluated by sliding the query across all possible rotational windows and computing sequence identity and similarity within each frame. The maximum score across all windows is retained. This procedure ensures that scoring reflects the rotational symmetry of cyclic peptides rather than the fixed orientation assumed in linear alignments.

**Figure 4.**
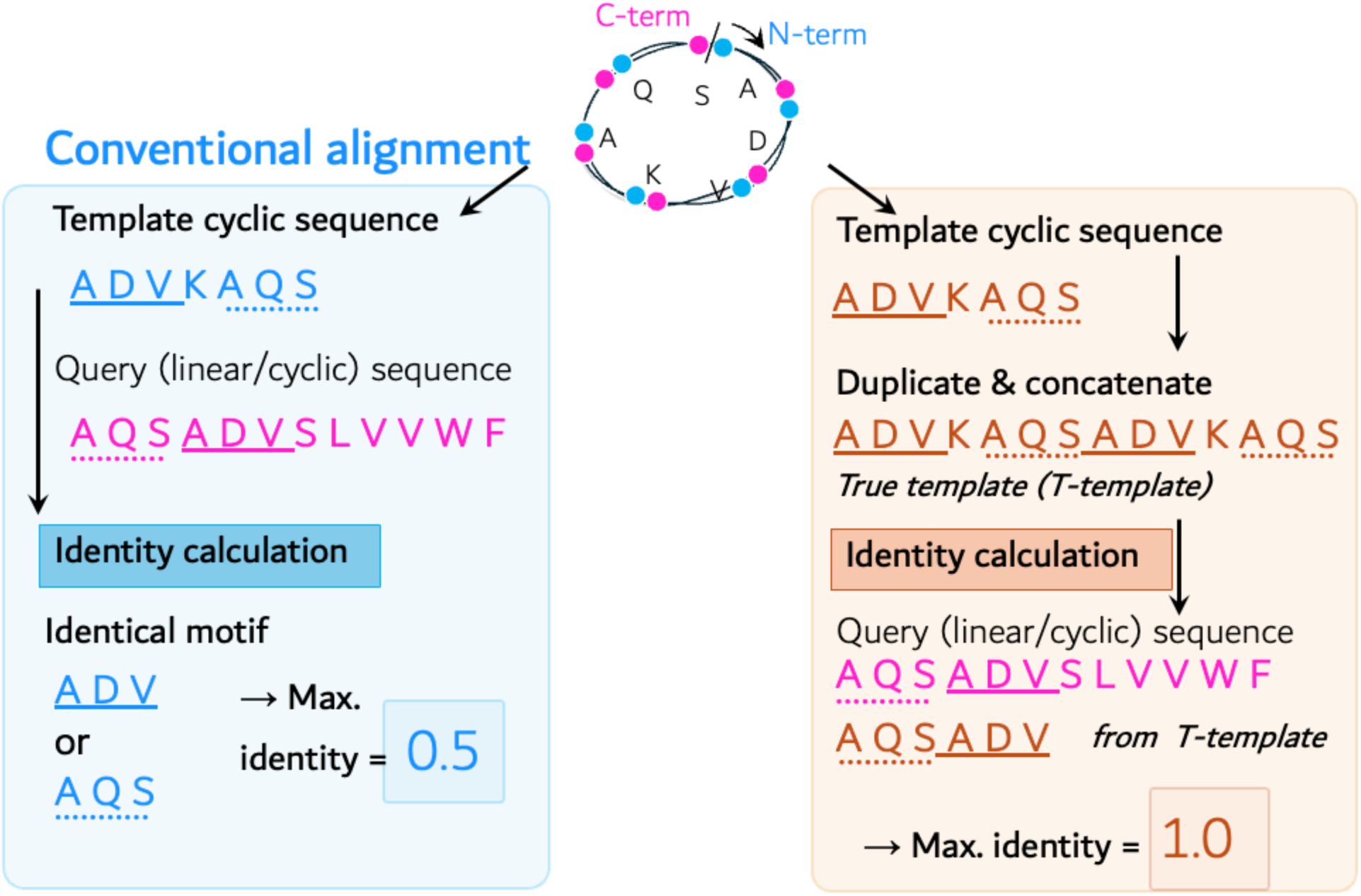
Comparison between conventional linear sequence alignment and cyclicity-aware alignment for cyclic peptides. In the conventional approach (left), sequence identity is computed using the linear template, which fails to account for the circular nature of cyclic peptides and results in partial motif matching and underestimated identity scores (maximum identity = 0.5). In contrast, the cyclicity-aware method (right) duplicates and concatenates the template sequence to generate a “true template” (T-template), enabling proper local alignment across the cyclization junction. This approach captures continuous motifs spanning the N- and C-termini, leading to accurate motif recovery and improved sequence identity (maximum identity =1).

**Figure 5.**
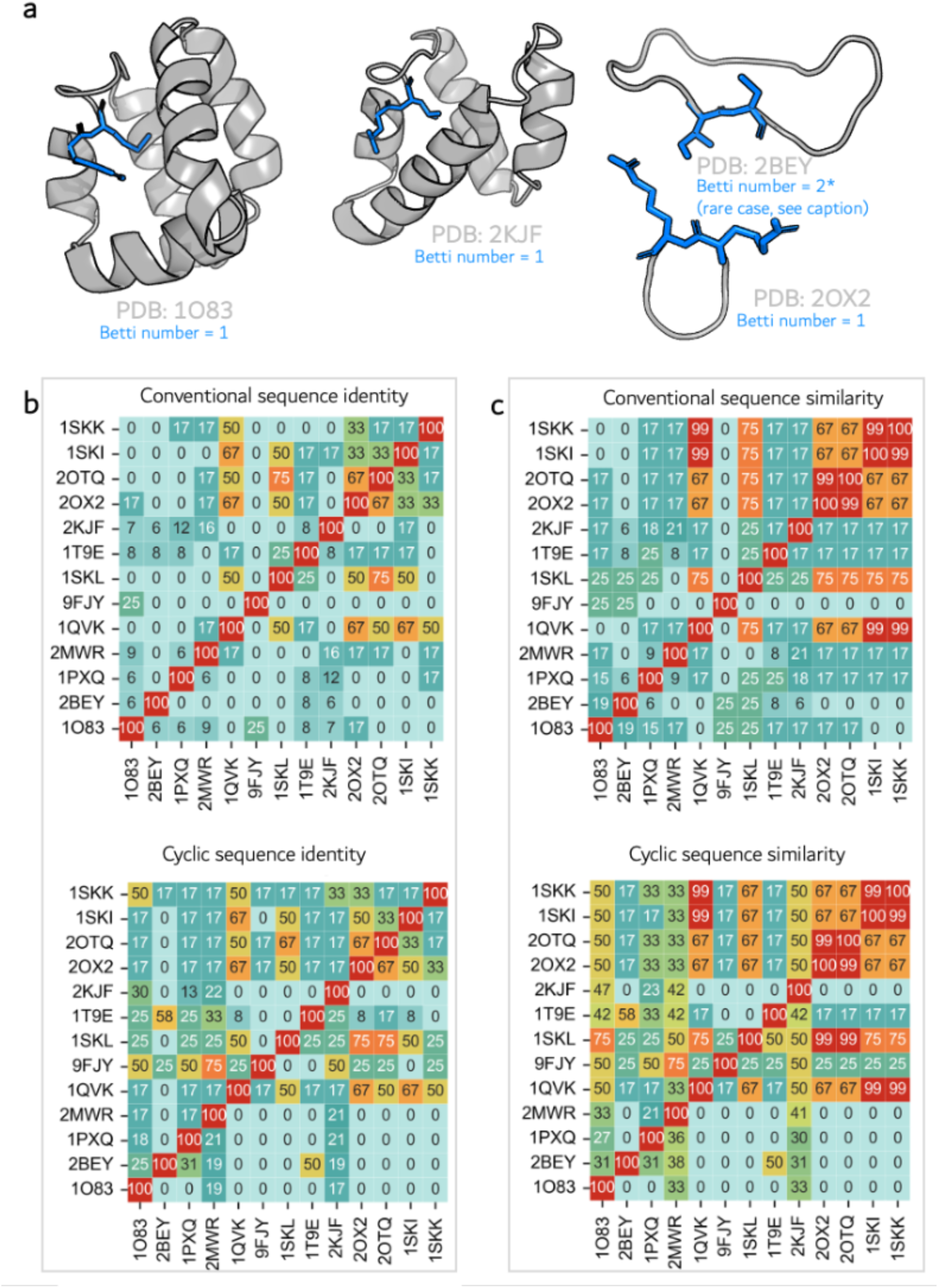
Conventional vs. cyclicity-aware sequence similarity for e2e cyclic peptides. **(a)** Representative structural diversity among e2e cyclized peptides in Cyclome930. 2BEY is a rare case where terminal Cys is involved in e2e cyclisation as well as e2s. For clarity in classifying, it as e2e, we highlighted only e2e linkage. First betti number, was able to identify two cycles sharing the same Cys *b*1(*G*) = 2 **(b)** Conventional sequence identity vs cyclic sequence identity and matrices. **(c)** Conventional sequence similarity vs cyclic sequence similarity matrices. Because conventional scoring compares peptides in a fixed N→C register, interchanging the query and template does not change the computed value, resulting in perfectly symmetric matrices in which the upper and lower triangles are identical. Cyclicity-aware identity and similarity matrices computed using the T-template rotational sliding protocol. Unlike conventional scoring, the cyclic protocol is orientation- and length-dependent: a query of length *L*₍q₎ is slid across a duplicated template of length 2·*L*₍t₎. This breaks symmetry and leads to pronounced differences between the upper and lower triangles. The cyclicity-aware method reveals biologically meaningful relationships that are obscured by conventional linear comparison.

**Figure 5b-c** illustrates the difference in conventional sequence identity/similarity vs cyclicity aware identity/similarity calculations. Because linear methods treat a peptide as a fixed N→C string and rely on positional comparison, interchanging the template and query sequences has no effect on the computed values. As a result, the upper and lower triangles of the matrices are perfectly symmetric, with the diagonal forming a clean hypotenuse. This symmetry reflects the commutative nature of linear similarity calculations, where similarity (A, B) = similarity (B, A). However, this mathematical convenience does not account for circular topology or rotational register matching.

In contrast, the cyclicity-aware identity and similarity matrices reveal a fundamentally different pattern. Because the T-template protocol evaluates the best rotational alignment of the template relative to the query, reversing the roles of template and query can yield different optimal matches, or no valid match at all. Cyclic peptides do not have fixed termini, and in our implementation, identity and similarity is evaluated by sliding a query of length L₍q₎ across a concatenated template of length 2×L₍t₎. This directional definition breaks the symmetry inherent to linear scoring and introduces a strong dependence on sequence length. When the template is much longer than the query, the duplicated template provides many possible windows, allowing local matches to be detected. In the opposite orientation, however, no valid window exists if the query length exceeds twice the template length (L₍q₎ > 2×L₍t₎), forcing the cyclic similarity and identity to 0% even when conventional linear methods report substantial similarity. For example, comparing a 6-residue peptide (PDB: 2OX2) against a 60-residue peptide (PDB: 2KJF) using 2KJF as the template corresponds to scanning a 6-residue query across a 120-residue T-template and yields 50% cyclic similarity. Reversing the roles using 2OX2 as the template and 2KJF as the query attempts to align a 60-residue query onto a duplicated 6-residue template with an effective length of 12 residues, which cannot produce any valid window and therefore yields 0% cyclic similarity. In contrast, conventional linear sequence similarity calculations report 17% similarity in both orientations. Consequently, cyclic identity and similarity matrices are inherently asymmetric, with the upper and lower triangles differing substantially. Several peptide pairs, such as 2OX2–2KJF, display pronounced asymmetry that reflects template–query definition and length-dependent constraints rather than a simple symmetric notion of sequence similarity.

Importantly, this orientation- and length-dependent behavior also reveals true cyclic motif conservation that linear scoring underestimates. For instance, the peptide pair 2OX2 (T-template) and 1SKL (query) exhibits 99% cyclic similarity, reflecting near-perfect alignment of a conserved cyclic motif. Conventional linear scoring reports only 75% similarity for this pair because it ignores rotational equivalence and enforces a fixed N→C register. Thus, the asymmetry observed in the cyclicity-aware similarity and identity calculations arises from the spatial pattern of residue arrangement imposed by cyclization topology. Breaking cyclic peptides into linear chains through conventional sequence comparison dilutes and often substantially underestimates their inherent similarity. Together, these observations demonstrate that cyclicity-aware similarity and identity provide a more faithful representation of structural relationships among cyclic peptides than conventional linear methods.

Single side-chain cyclization, as observed in side-to-side (s2s) and side-to-end (s2e) peptides, introduces a covalent constraint while preserving the linear residue register of the backbone. Unlike head-to-tail (e2e) cyclization, which rotates the entire sequence and disrupts linear correspondence, s2s and s2e cyclization geometries maintain a one-to-one mapping between linear sequence positions and their spatial counterparts. Consequently, conventional linear sequence identity and similarity calculations remain meaningful for these classes of cyclic peptides because the relevant functional and structural motifs retain their order along the backbone. **Figure6a** and **6b** illustrate representative structures of s2s and s2e peptides, respectively. The cyclizing residues (shown in blue) impose local constraints but do not reorder the positions of the remaining residues. As a result, backbone topology is preserved even though the peptides adopt compact three-dimensional folds. This preservation of linear register explains why linear alignment captures true structural similarity for s2s and s2e peptides, in contrast to e2e peptides where circular permutation disrupts positional correspondence. The matrices of sequence identity **(Figure 6c, e)** and similarity **(Figure 6d, f)** exhibit symmetric patterns, an outcome expected only when residue correspondence is unambiguous, as in linear peptides. Off-diagonal values reveal gradients of relatedness and clusters of peptides sharing moderate to high identity or similarity. Similarity matrices further highlight chemically conserved features even when strict identity is low, reinforcing that preserved linear ordering aligns residues occupying equivalent structural positions. Together, the structural representations and heatmaps show that for s2s and s2e peptides, conventional linear identity and similarity measurements capture true structural similarity, in contrast to e2e peptides where linearization disrupts positional mapping and obscures biologically relevant relationships. However, when peptides contain multiple s2s or s2e cyclization points, the backbone is partitioned into several interconnected loops, and linear sequence identity and similarity no longer reflect spatial residue arrangement. In such cases, residues that interact in three dimensions may be distant in sequence, and residues adjacent in sequence may be spatially separated.

**Figure 6.**
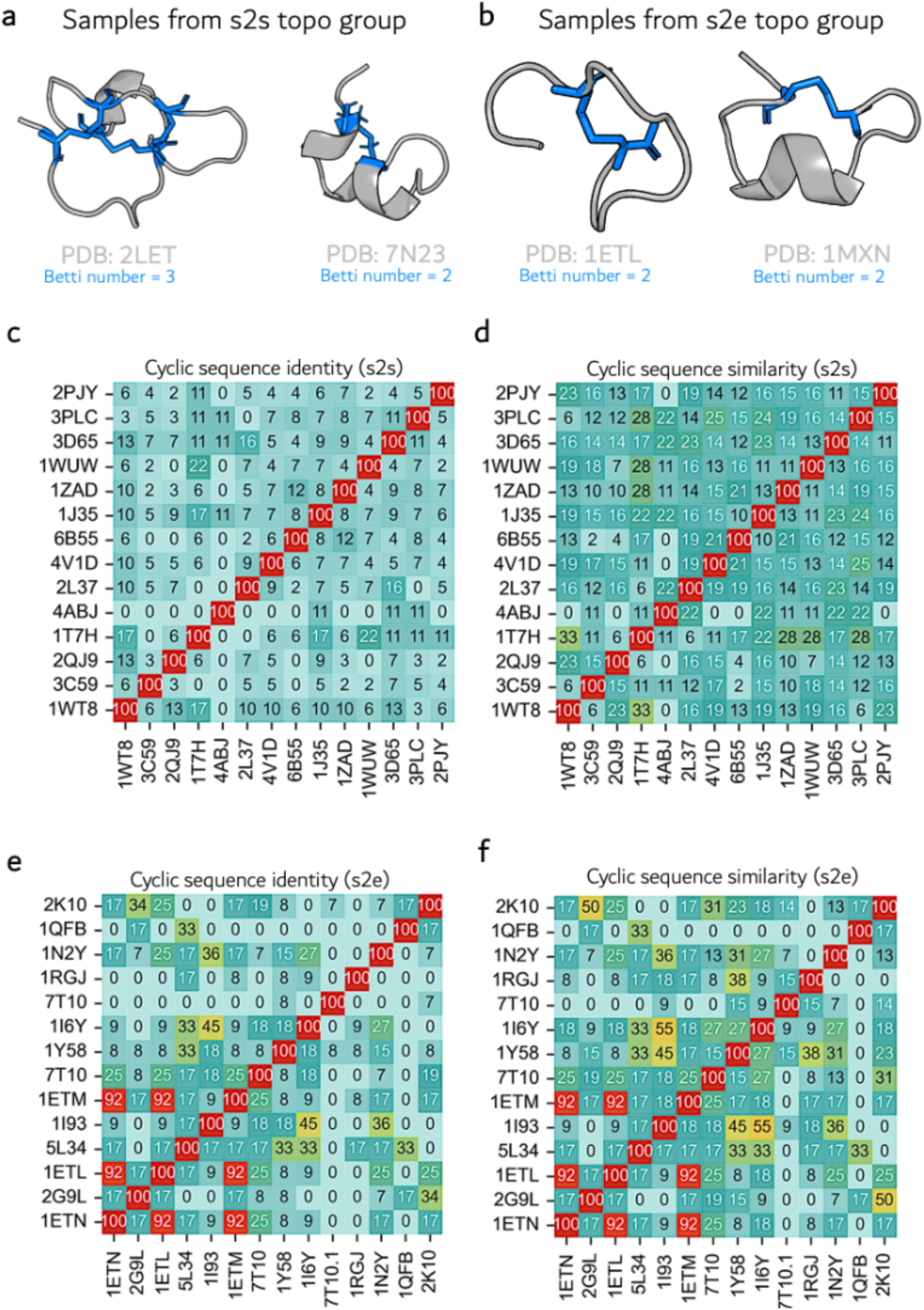
Structural and sequence relationships in s2s and s2e cyclic peptides. (**a–b**) Representative structures of side-to-side (s2s) and side-to-end (s2e) peptides illustrating that a single side-chain cyclization (blue) constrains the backbone without disrupting the linear residue register. Because residue order is preserved, linear sequence alignment accurately reflects spatial correspondence. (**c–f**) Pairwise sequence identity **(c, e**) and similarity (**d, f**) matrices for s2s and s2e peptide sets. The symmetric heatmaps show that linear scoring captures meaningful relationships, including clustering of peptides with conserved spatial residue patterns. In contrast to head-to-tail cyclic peptides, single side-chain cyclization preserves positional mapping along the backbone. However, when peptides contain multiple s2s or s2e cyclization points, the backbone is partitioned into several interconnected loops, and linear sequence identity/similarity no longer reflects spatial residue arrangement; in such cases, residues that interact in 3D may be distant in sequence and vice versa.

Cyclic peptides containing both e2e and s2s (e2e+s2s), **Figure 7a** also present a challenge for sequence-based comparison because their linear sequences do not reflect true covalent connectivity through the backbone and side chains. To quantify the effect of topology on sequence relationships, we computed pairwise similarity and identity using both conventional linear sequences and a cyclicity-aware T-template representation that explicitly encodes the residue adjacency produced by e2e+s2s cyclization (see ***Cyclicity-aware -sequence Similarity Calculator in Methods*** for details). When conventional linear sequences were compared, the resulting similarity matrix displayed a diffuse pattern dominated by low pairwise similarity values (**Figure 7c**). This occurs because linear alignment assumes a continuous N→C backbone and ignores the reshuffling of spatial positions and new adjacency patterns introduced by cyclization. As a result, peptides that have similar sequence and structure but different in the placement of cyclization points appear weakly related or unrelated under linear comparison. In contrast, the T-template–based cyclicity-aware similarity matrix revealed distinct blocks of higher similarity (**Figure 7d**). By reorganizing each sequence to reflect the actual connectivity established by e2e and s2s bonds, the method aligns structurally equivalent residues across peptides and uncovers higher-order similarity signals that are obscured in linear space. A similar trend was observed for pairwise sequence identity (**S. Figure 1**), where the conventional identity matrix was dominated by uniformly low values with few discernible clusters. Many peptides shared less than 10% apparent identity despite belonging to related structural families. When identity was recalculated using cyclicity-aware T-templates, several coherent regions of increased identity emerged. Peptides sharing similar ring sizes, cyclization patterns, or conserved structural motifs, but differing in linear residue order due to variable s2s placements showed substantially higher identity once residue ordering was corrected to match macrocycle topology. Thus, cyclicity-aware identity successfully recovers relationships that are invisible under conventional analysis.

**Figure 7.**
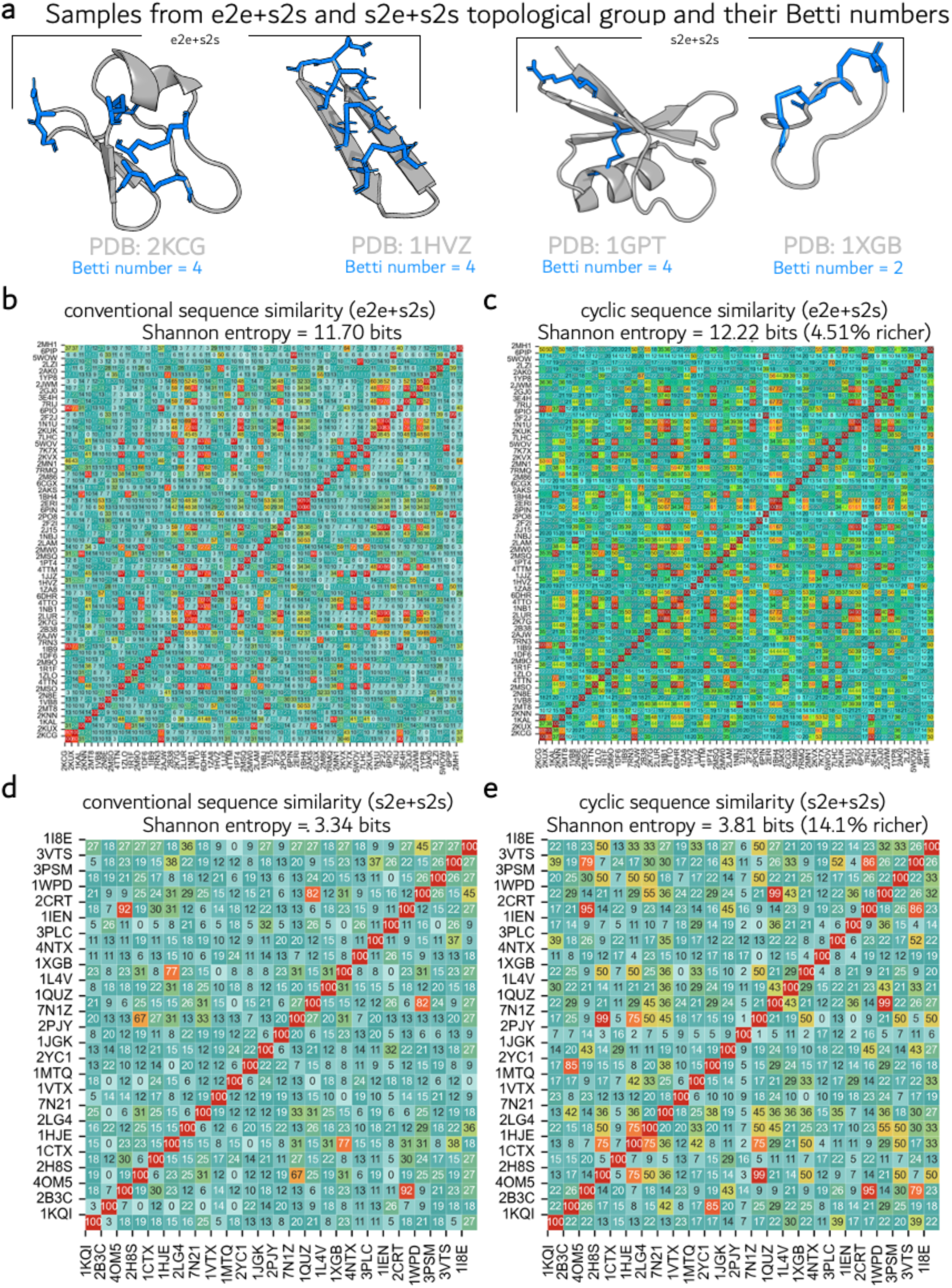
Sequence similarity comparison of e2e+s2s and s2e+s2s cyclic peptides using conventional and cyclization-aware methods. **(a)** Representative structures of peptides containing both end-to-end and side-to-side (e2e+s2s) and side-to-end with side-to-side (s2e+s2s) cyclization, illustrating how multiple covalent linkages reorganize backbone connectivity. **(b, d)** Conventional sequence similarity matrix computed directly from the linear amino-acid sequences. Because this approach does not account for ring closures or the altered residue adjacency imposed by e2e+s2s and s2e+s2s cyclization, many peptides appear weakly related despite similar folded topologies. **(c, e)** Cyclicity-aware sequence similarity matrix computed using the T-template framework, in which sequence rearrangements reflect the true covalent connectivity of e2e+s2s and s2e+s2s cyclic peptides. Incorporating cyclization information reveals additional similarity relationships that remain hidden under conventional linear comparison.

Following the similar trend, for cyclic peptides containing combined side-to-end and side-to-side (s2e+s2s) cyclization (**Figure 7a**), conventional linear sequence identity and similarity calculations fail to capture the true relationship between sequence and structure. As shown in the conventional similarity matrix (**Figure 7d**), several peptide pairs exhibit low to moderate similarity despite sharing conserved cyclization-constrained residue arrangements. As mentioned earlier, the discrepancy arises because linear alignment treats the sequence as an open chain and does not account for the enforced proximity of residues introduced by multiple cyclization linkages. In contrast, the cyclicity-aware protocol produces markedly higher similarity patterns (14.1 % higher Shannon entropy), with enhanced clustering of peptides that share common spatial residue motifs even when their linear register differs (**Figure 7e**). Notably, peptide pairs that appear weakly related under linear similarity often show substantially higher cyclic sequence similarity, reflecting preservation of three-dimensional residue organization rather than linear order. These results demonstrate that, unlike simple s2s or s2e peptides where linear similarity can approximate spatial correspondence, the presence of multiple cyclization constraints disrupts the linear–spatial mapping. Consequently, linear similarity systematically underestimates relatedness for s2e+s2s peptides, whereas cyclicity-aware calculations more accurately reflect their underlying structural and topological similarity.

Across both similarity and identity metrics (**Figure 5**, **Figure 7**, **S. Figure 1, S. Figure 2**), the results consistently show that conventional sequence representations fail to capture meaningful relationships among e2e, e2e+s2s and s2e+s2s classes of cyclic peptides. Incorporating cyclization topology via T-templates (i) restores correspondence between residues that are functionally or spatially aligned, (ii) reveals clusters of related peptides that linear methods miss, and (iii) provides a more accurate basis for annotation, clustering, and rational design of structurally rich macrocycles. Together, these findings demonstrate that recognizing cyclic topology is essential for sequence-based analysis of multiply cyclized peptides and that cyclicity-aware representations substantially improve the interpretability and biological relevance of peptide similarity metrics.

### REMD-derived structural priors for extracting melting points of cyclic peptides

Replica-exchange molecular dynamics (REMD) simulations were employed as an enhanced sampling strategy to extensively explore the conformational landscapes of cyclic peptides across a wide temperature range (**Figure 8a**). We leveraged the temperature-dependent structural ensembles generated by REMD to identify melting-like transitions based on coordinated changes in conformational sampling (Ramachandran angles), compactness (radius of gyration), backbone flexibility (RMSF) and deviation of overall structure from the starting conformer (RMSD).

**Figure 8.**
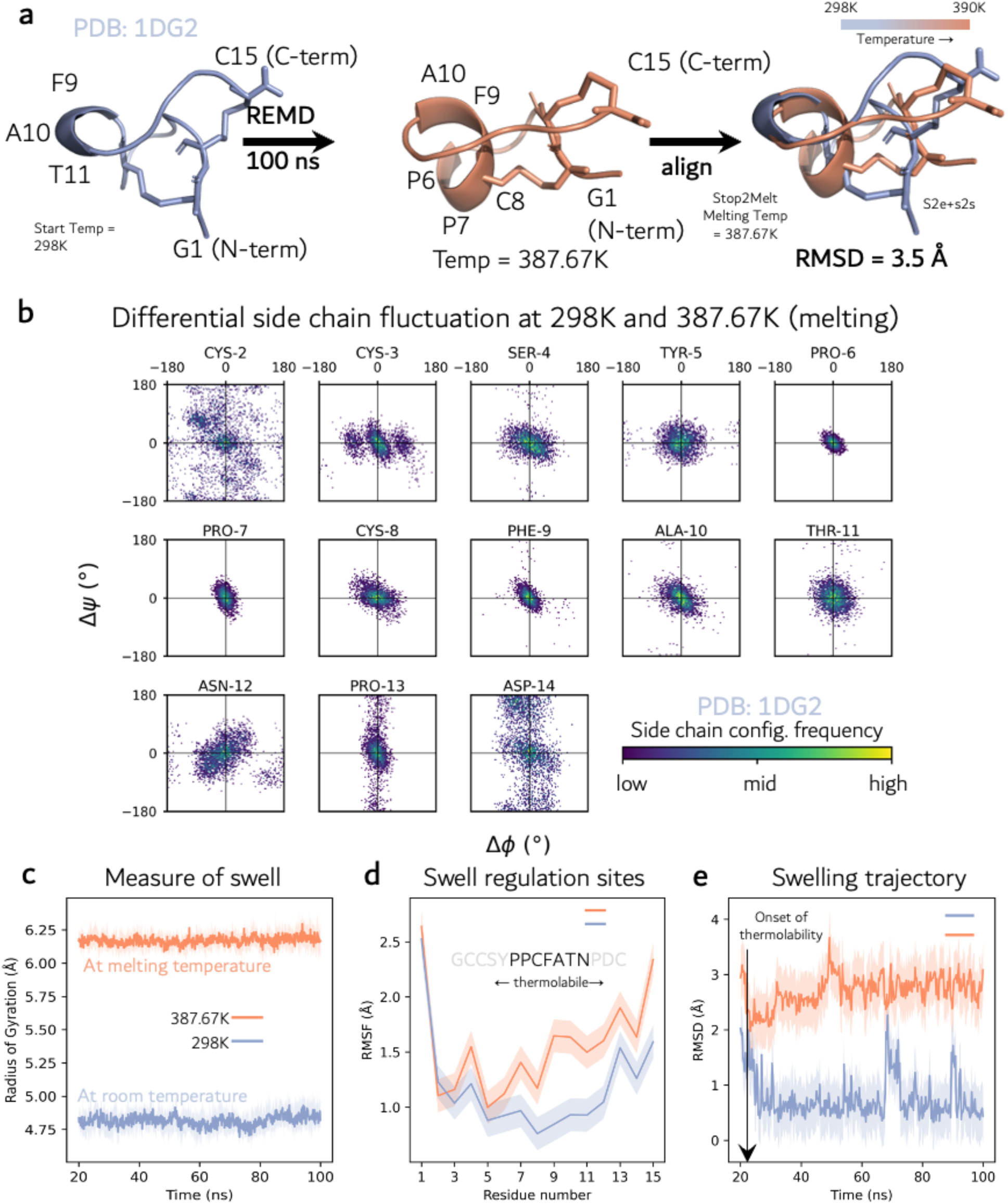
REMD-derived structural indicators for identifying melting-like transitions in cyclic peptides (STop2Melt). **(a)** Changes in cyclic peptide structure indicating melting-like transitions (**b**) Ramachandran difference maps (Δφ–Δψ) highlighting changes in backbone dihedral occupancy between ensembles below and above the inferred melting point (STop2Melt). The non-overlapping dihedral populations indicate a sharp conformational transition. **(c)** Radius of gyration (Rg) as a function of temperature, showing a clear transition at STop2Melt = 387.67 K. Structures above STop2Melt consistently exhibit Rg > 6 Å, whereas structures below STop2Melt remain compact with Rg < 5 Å, with minimal overlap between regimes. **(d)** Residue-wise root-mean-square fluctuation (RMSF) profiles, demonstrating uniformly low fluctuations below STop2Melt and pronounced increases at and above STop2Melt, particularly for residues 6-12, identifying localized hotspots of thermal destabilization. **(e)** Root-mean-square deviation (RMSD) time series relative to the reference structure, showing stable, narrowly distributed values below STop2Melt and broader, elevated RMSD values at and above STop2Melt, consistent with increased conformational diversity without complete unfolding. Together, these REMD-derived structural descriptors—dihedral dispersion, Rg expansion, and RMSF/RMSD broadening—provide a coherent set of indicators for identifying melting-like transitions in cyclic peptides and for extracting inferred melting points (STop2Melt).

We tracked residue-wise backbone conformational sampling through dihedral angles (Ramachandran angles) (**S. Figure 3a, b**) and calculated the difference map (Δφ, Δψ) from the simulation trajectory below and at/above the melting point, STop2Melt (**Figure 8b**). Below the inferred melting point, cyclic peptides populate narrow and well-defined backbone dihedral regions (**S. Figure 3a**), reflecting strong topological constraints imposed by cyclization. Above the inferred melting point, several residues exhibit markedly broadened dihedral distributions (**S. Figure 3b**).

Global structural organization, quantified by the radius of gyration, Rg (**Figure 8c**), exhibits a sharp transition at the inferred melting point (STop2Melt = 387.67 K for 1DG2), an indication of peptide swelling/denaturation. Above STop2Melt, the radius of gyration consistently exceeds 6 Å, indicating partial expansion/swelling of the cyclic scaffold, whereas below STop2Melt, Rg remains compact and below 5 Å. The near absence of overlap between these Rg regimes provides a strong structural signature for the melting-like transition.

Residue-wise RMSF profiles (**Figure 8d**) further delineate this transition. Structures below the melting point exhibit uniformly low fluctuations, while at and above the melting point, pronounced increases in RMSF are observed for residues Pro6, Pro7, Cys8, Phe9, Ala10, Thr11, Asn12 identifying these positions as local hotspots of thermal destabilization/ swell regulation sites. The selective nature of these fluctuations supports a localized, noncooperative melting mechanism rather than uniform backbone disorder.

RMSD time series (**Figure 8e**) show behavior consistent with both Rg and RMSF trends. Below STop2Melt, RMSD values remain low and narrowly distributed, reflecting structural stability. At and above STop2Melt, RMSD values increase and broaden, indicating access to alternative conformational basins. As with Rg and RMSF, RMSD distributions above the melting point are distinct and minimally overlapping with those below, reinforcing the presence of a well-defined transition.

Using the concurrent onset of dihedral dispersion, Rg expansion from < 5Å to > 6Å, and residue-specific RMSF/RMSD broadening as structural indicators, we identified a characteristic transition temperature for each peptide, as its inferred melting point (STop2Melt). This procedure was applied systematically to all peptides in Cyclome930, yielding a comprehensive set of STop2Melt annotations. The extracted STop2Melt values for all peptides are reported in **Supplementary Data 1**.

Importantly, the REMD-derived STop2Melt values are consistent with experimental observations for well-characterized cyclic peptides. For the prototypical cyclotide Kalata B1 (PDB: 4TTN), our calculations yield an STop2Melt of 382.38 K. This is in strong agreement with experimental studies by Colgrave and Craik, which showed that Kalata B1 retains its structural integrity upon heating from 310 K to 370 K, even as temperatures approach the boiling point of water.^24^ This concordance supports the physical validity of our transition-based definition of melting and highlights the ability of our framework to capture the exceptional thermal resilience characteristic of cyclic peptide scaffolds.

Together, these results demonstrate that REMD, while primarily an enhanced sampling method enables robust identification of melting-like transitions in cyclic peptides when interpreted through ensemble-resolved structural descriptors. The clear, non-overlapping separation between structural regimes below and above STop2Melt provides strong internal consistency for the inferred melting points and establishes a physically grounded stability reference for large-scale analysis of cyclic peptide thermostability.

The REMD-derived STop2Melt values described above provide a physically motivated stability reference for cyclic peptides, capturing the onset of melting-like transitions through coordinated changes in backbone conformational sampling, compactness, and flexibility. While REMD enables robust identification of these transitions, its computational expense precludes routine application at scale. We therefore leveraged the REMD-inferred STop2Melt annotations for Cyclome930 as supervised targets to develop a data-driven predictor capable of estimating melting-point priors directly from peptide sequence and cyclization topology.

### STop2Melt predictor: Machine-learning model for melting-point prediction of cyclic peptides

We trained a supervised regression model to predict STop2Melt, for cyclic peptides in the Cyclome930 dataset. Nine regression models were trained and evaluated using cluster-aware train, validation, and test splits, allowing a rigorous assessment of generalization across distinct cyclic peptide scaffolds (**Figure 9a**). All models used identical splits and comparable hyperparameter settings, enabling direct comparison of feature representations. Our analysis highlights the importance of cyclicity-aware embeddings for machine-learning tasks involving cyclic peptides. Models trained using ESMc sequence embeddings alone showed poor predictive performance for STop2Melt (R^2^≤ 2.8), indicating that linear sequence representations are insufficient to capture the dominant determinants of cyclic peptide stability.

**Figure 9.**
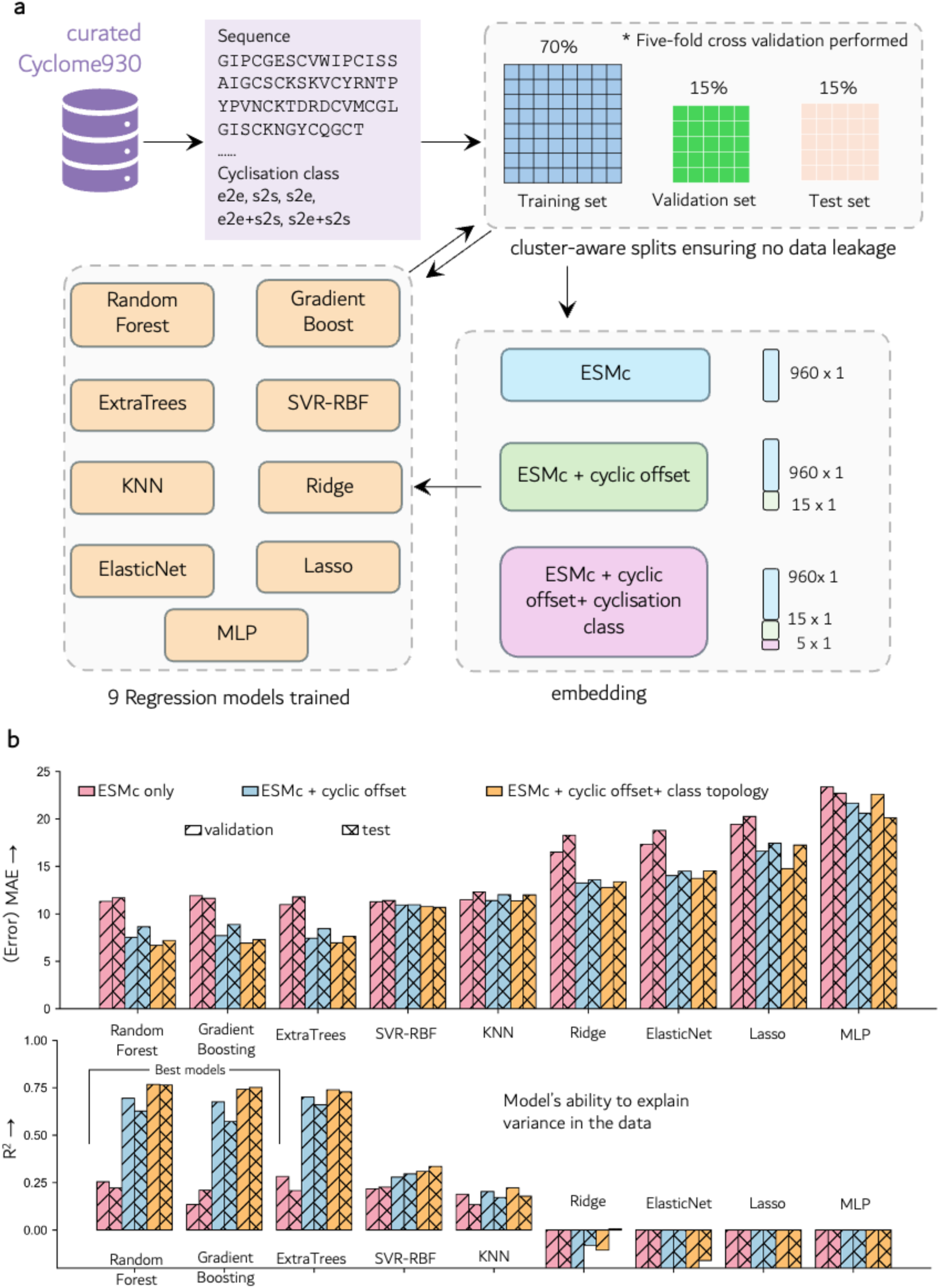
The architecture of STop2Melt for melting point prediction, and its performance under different embedding strategies. **(a)** The workflow of STop2Melt organizes Cyclome930 into clusters based on shared sequence and cyclization connectivity and splits them in a cluster-awar**e** manner into training, validation, and test sets to prevent information leakage. Peptide sequences are embedded using ESMc, with three feature settings evaluated: ESMc embeddings alone, ESMc embeddings augmented with cyclic offset features, and ESMc embeddings augmented with both cyclic offsets and cyclization-class information. These embeddings are used to train multiple regression models, including tree-based, kernel-based, linear, instance-based, and neural network regressors. The trained models predict STop2Melt, a REMD-derived melting-point proxy, for novel cyclic peptides, with results summarized across all models and the most confident prediction highlighted. (**b**) Performance of STop2Melt in terms of Mean absolute error (MAE) and coefficient of determination (R^2^) for nine regression models evaluated using cluster-aware test splits. The results are shown for three feature settings: ESMc embeddings only, ESMc embeddings augmented with cyclic offsets, and ESMc embeddings augmented with cyclic offsets and cyclization-class topology descriptors. Bars are grouped by regression model, with validation and test performance distinguished by hatch patterns. Incorporation of cyclic offsets leads to substantial reductions in MAE and R^2^ increases across ensemble models, while the addition of topology descriptors provides further gains for the best-performing methods, highlighting the importance of cyclicity-aware representations for predicting cyclic peptide melting behavior.

To systematically assess the contribution of cyclic topology, we compared three feature settings: (i) ESMc embeddings only, (ii) ESMc embeddings concatenated with cyclic offset and cyclization-class topology descriptors, and (iii) topology-ablated control models using ESMc embeddings with cyclic offsets but without higher-level topology descriptors.

All comparisons were performed using identical data splits and model configurations. Cyclic offsets encode effective shortest-path distances between residues under cyclic connectivity and provide a compact, sequence-independent representation of cyclization-induced constraints. These offsets complement ESMc embeddings by explicitly capturing residue adjacency relationships that are not reflected in linear sequence order. By comparing ESMc only models with cyclic-offset–augmented models, we directly evaluated the impact of employing cyclicity aware embeddings on melting-point prediction.

### ESMc embeddings alone are insufficient for STop2Melt prediction

Models trained using ESMc sequence embeddings alone exhibited limited predictive performance across all regression algorithms (**Figure 9b**). Test R^2^ values remained low, 0.23 for the best performing model (**S. Table 1, S. Figure 4a**), with corresponding MAE values of approximately 11 K for the best-performing ensemble models (Random Forest and ExtraTrees). Linear models and the multilayer perceptron performed particularly poorly under cluster-aware evaluation, yielding strongly negative R^2^ values and high MAE. These results indicate that linear sequence-derived representations, even when learned from protein large language models, are insufficient to capture the dominant determinants of cyclic peptide melting behavior. In particular, ESMc embeddings do not explicitly encode the topological constraints imposed by cyclization, which strongly influence peptide stability.

### Cyclic offsets encode essential connectivity information

To explicitly incorporate cyclicity, we augmented ESMc embeddings with cyclic offset descriptors, which encode effective shortest-path distances between residues under cyclic connectivity. These offsets provide a compact, sequence-independent representation of residue adjacency along the cyclic backbone and across cyclization links. Incorporation of cyclic offsets led to pronounced and systematic improvements in predictive performance across nearly all models (**Figure 9b**). Tree-based ensemble methods showed the largest gains, with test R^2^ increasing to 0.66 and MAE decreasing substantially relative to ESMc only baselines (11 K to 8 K). Importantly, validation and test performance tracked closely, indicating improved generalization rather than overfitting (**Figure 9b, S. Table 2, S. Figure 4b**). These results demonstrate that explicit encoding of cyclic connectivity is essential for modeling STop2Melt and that cyclic offsets capture structural constraints that are not reflected in linear sequence order.

### Topology-aware descriptors further improve performance

While cyclic offsets encode local residue adjacency, they do not capture higher-level architectural properties such as global connectivity patterns or cyclization class identity. To address this, we incorporated cyclization-class topology descriptors as additional global features.

Models trained with ESMc embeddings, cyclic offsets, and topology descriptors consistently achieved the best overall performance (**Figure 9b, S. Table 3, S. Figure 4c**). For the top-performing ensemble models, test R^2^ values increased further, reaching 0.76, with corresponding reductions in MAE (7.2 K). These gains were observed consistently across validation and test sets, confirming that topology-aware features provide complementary information beyond cyclic offsets alone.

### Ensemble models capture nonlinear cyclicity–stability relationships

Across all feature settings, tree-based ensemble models (Random Forest, Gradient Boosting, and ExtraTrees) outperformed linear, kernel-based, and neural network models (**Figure 9a, b**). This trend highlights the importance of modeling nonlinear interactions between sequence-derived features and cyclic topology. In contrast, linear models and the MLP failed to leverage cyclicity-aware descriptors effectively, underscoring the complexity of the underlying stability landscape.

### Structural interpretation and mechanistic consistency

The dominant contribution of cyclic offsets is consistent with REMD-derived structural signatures of melting, which indicate that STop2Melt corresponds to a topologically constrained transition involving changes in cyclic scaffold compactness, selective residue-level flexibility, and altered residue–residue proximity. These effects are inherently invisible to linear sequence embeddings but are directly reflected in cyclic-offset representations.

Together, these results establish cyclic offsets as a critical inductive bias for machine-learning models of cyclic peptide stability. By bridging REMD-resolved structural transitions with scalable, sequence-based representations, the STop2Melt predictor provides a physically grounded and generalizable framework for melting-point prediction of cyclic peptides.

### CritiCL (a classifier model for peptides binding to critical minerals) screens Cyclome930 for critical mineral capture

To identify peptide sequences with potential affinity toward critical mineral ions, we performed zero-short prompting for Cyclome930 on CritiCL, a multiclass machine learning classifier developed for classifying proteins and peptides based on the metal ion they are capable of binding, distinguishing proteins and peptides based on metal-binding specificity among lanthanides (Ln^3+^), cobalt (Co^2+^), nickel (Ni^2+^), and manganese (Mn^2+^) and other transition metal ions (**Figure 10**). Ensemble tree-based models, particularly Random Forest and XGBoost, yielded the most stable performance across evaluation metrics, whereas neural network and support vector machine models showed larger variability in predictions across folds (**Table 2**). Overall, the models achieved high classification accuracy and balanced performance across the five metal classes, suggesting that the sequence embeddings retained meaningful information related to metal-binding specificity. We enabled cyclicity awareness for CritiCL by incorporating cyclic offsets/linear offsets and cyclic/ linear topology descriptors as described for STop2Melt predictor.

**Table 2.** Performance metrics for all the four classification models (XGBoost, Random Forest, Support Vector Machine and Multi-Layer Perceptron) trained within CritiCL.

| Model | CV used | Accuracy | Macro F1 | Weighted F1 | MCC | Macro ROC-AUC |
| --- | --- | --- | --- | --- | --- | --- |
| **XGB** | StratifiedKFold(n_splits=5) | 0.79 | 0.81 | 0.79 | 0.72 | 0.96 |
| **RF** | StratifiedKFold(n_splits=5) | 0.78 | 0.80 | 0.78 | 0.71 | 0.95 |
| **MLP** | StratifiedKFold(n_splits=5) | 0.75 | 0.77 | 0.75 | 0.67 | 0.92 |
| **SVM** | StratifiedKFold(n_splits=5) | 0.75 | 0.77 | 0.75 | 0.66 | 0.93 |

**Figure 10.**
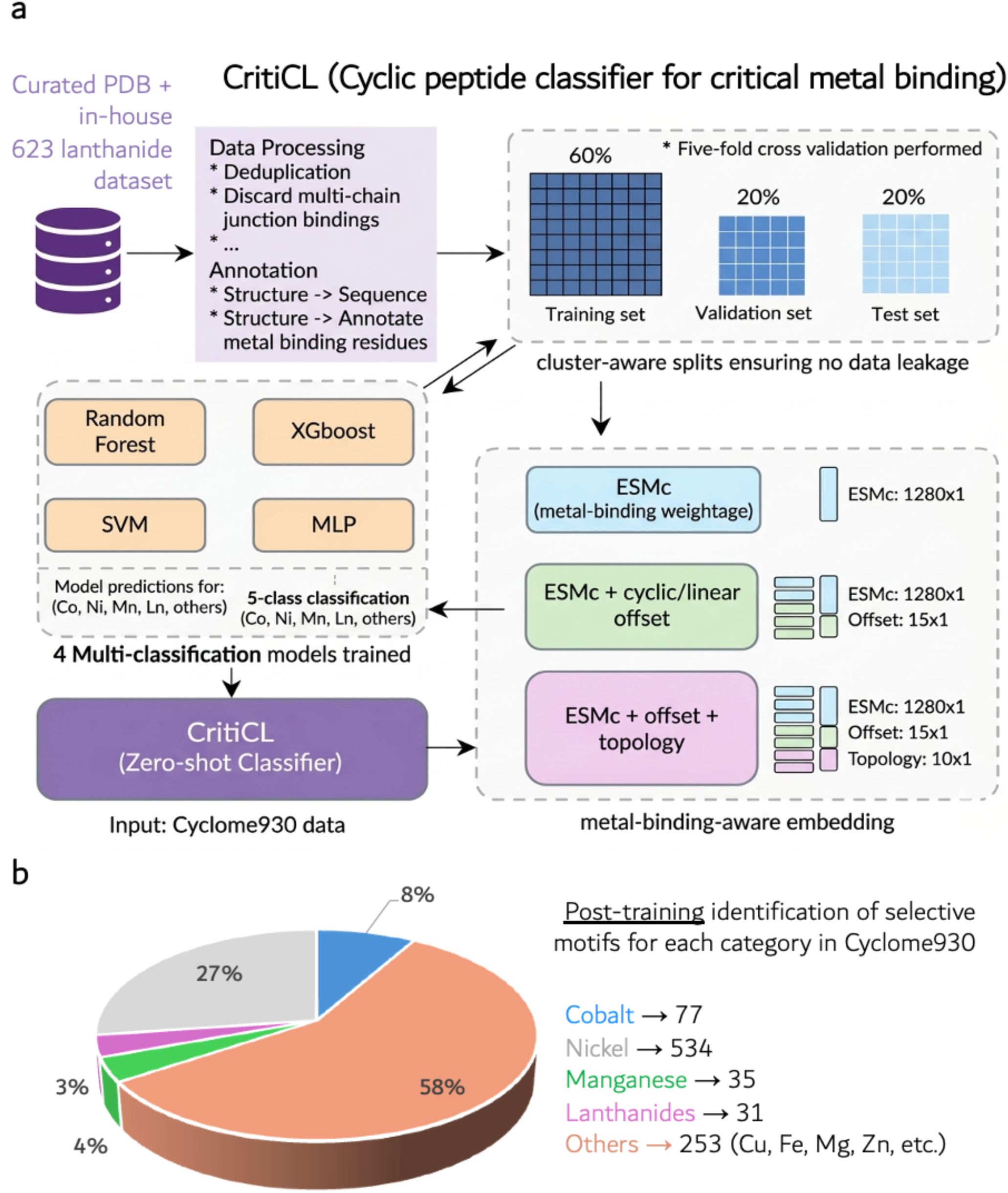
Overview of the CritiCL framework for cyclic peptide classification targeting critical metal binding. (**a**) A curated metal binding PDBs along with in-house 623 lanthanide-binding protein structures was processed through deduplication and filtering to remove multi-chain junction artifacts, followed by annotation of metal-binding residues from structure-to-sequence mapping. Cluster-aware data splitting (60% training, 20% validation, 20% test) with five-fold cross-validation ensured no data leakage. Multiple classifiers (Random Forest, XGBoost, SVM and MLP) were trained for five-class prediction (Co^2+^, Ni ^2+^, Mn ^2+^, Ln ^3+^, others). Protein sequences were encoded using metal-binding–aware embeddings derived from ESMc, augmented with cyclic/linear offsets and topological features to capture structural context. The best-performing models were integrated into CritiCL, enabling zero-shot classification of cyclic peptides (Cyclome930) for selective metal binding prediction. (**b**) Results from zero-shot classification of Cyclome930 showing the predicted distribution of metal-binding classes (Co, Ni, Mn, Ln, and others).

**Figure 10.** Architecture for CritiCL (Cyclic Peptide Classifier for critical metal binding). A curated metal binding PDBs along with in-house 623 lanthanide-binding protein structures was processed through deduplication and filtering to remove multi-chain junction artifacts, followed by annotation of metal-binding residues from structure-to-sequence mapping. Cluster-aware data splitting (60% training, 20% validation, 20% test) with five-fold cross-validation ensured no data leakage. Multiple classifiers (Random Forest, XGBoost, SVM and MLP) were trained for five-class prediction (Co^2+^, Ni ^2+^, Mn ^2+^, Ln ^3+^, others). Protein sequences were encoded using metal-binding–aware embeddings derived from ESMc, augmented with cyclic/linear offsets and topological features to capture structural context. The best-performing models were integrated into CritiCL, enabling zero-shot classification of cyclic peptides (Cyclome930) for selective metal binding prediction. (b)

### Prediction confidence and metal-specific probability distributions

For each peptide sequence in Cyclome930, CritiCL produced probability scores corresponding to the five metal-binding classes. These scores provided a quantitative measure of prediction confidence and enabled ranking of candidate peptides according to their predicted binding preference. The probability distributions revealed that many cyclic peptides exhibited strong class separation (**Supplementary Data 2**), with a dominant probability assigned to a single metal ion class. In particular, several sequences displayed high confidence for Co^2+^ binding while maintaining substantially lower predicted probabilities for transition metal classes, indicating potential selectivity for rare-earth elements.

Based on the CritiCL confidence scores, Cyclome930 was ranked according to their predicted metal binding propensity (**Supplementary Data 2**). The peptides were classified into all the five classes, highlighting the varied potential of Cyclome930 in binding to metal binding and capture (**Figure 10b)**

## Discussion

Cyclic peptides are versatile biomolecules that bridge the gap between small molecules and proteins, offering enhanced stability, conformational rigidity, and functional diversity. Despite their growing importance in therapeutic and materials applications, systematic computational approaches for analyzing and predicting their thermal stability have remained limited. In this work, we address this gap by integrating large-scale data curation, topology-aware sequence analysis, physics-based molecular simulations, and machine-learning models specifically designed for cyclic peptides.

A central contribution of this study is the development of Cyclome930, a curated database of 930 unique cyclic peptides assembled from diverse public repositories. By substantially expanding previous datasets, Cyclome930 captures a broad range of sequence lengths, structural motifs, and biological origins. The integration of structural, biochemical, and biophysical annotations enables analyses that go beyond sequence alone, which is particularly important for cyclic peptides, where subtle differences in topology or crosslinking can strongly influence stability and function. As such, Cyclome930 provides both a robust foundation for model development and a valuable resource for the peptide research community.

Our results highlight the limitations of conventional linear sequence alignment methods for cyclic peptides. Because linear representations impose artificial termini, they often underestimate similarity and obscure conserved motifs. The cyclicity-aware sequence similarity framework introduced here explicitly accounts for rotational symmetry and complex cyclization patterns, enabling more faithful identification of related peptides. This capability is critical not only for comparative sequence analysis but also for machine-learning workflows, where similarity-aware data splitting is necessary to ensure realistic model evaluation.

REMD simulations provide a key link between molecular physics and data-driven modeling in this study. By sampling peptide conformations across a wide temperature range, REMD yields structure-based indicators of folding and unfolding behavior for cyclic peptides. Although absolute melting points are not always directly accessible, these simulation-derived stability metrics capture meaningful thermodynamic trends and correlate with experimental measurements, supporting their use as physically grounded training targets.

Building on these stability metrics, we show that machine-learning models incorporating cyclicity-aware embeddings can accurately predict melting like transitions in cyclic peptides. The combination of ESMc sequence embeddings with a cyclic offset enables the model to capture both sequence-level patterns and structural constraints imposed by cyclization. Models that ignore cyclic topology consistently underperform, underscoring the need for representations tailored to circular peptides.

From a practical perspective, the ability to predict thermal stability *in silico* has direct implications for cyclic peptide design and optimization. The STop2Melt framework presented here enables rapid screening of candidate peptides prior to synthesis, reducing experimental cost and guiding rational design. Although Cyclome930 represents only a fraction of the possible cyclic peptide sequence space and REMD simulations remain computationally demanding, continued data growth and methodological advances are expected to further improve predictive performance.

Overall, this work demonstrates that accurate prediction of cyclic peptide thermal stability requires cyclization-aware data resources and modeling strategies. By unifying curated databases, topology-aware sequence analysis, physics-based simulations, and machine learning, we establish a generalizable framework for understanding and engineering cyclic peptides, and we anticipate that these resources and methods will accelerate the discovery of stable cyclic peptide scaffolds.

Finally, we screened Cyclome930 for critical mineral binding and capture using a classification model (CritiCL) which is compatible to cyclicity aware embeddings for Cyclome930 and conventional embeddings for linear proteins and peptides. This work is a comprehensive work on cyclic peptides starting from data curation, sequence analysis, thermal stability prediction and screening them for critical mineral capture.

## Methods

### Compilation and Curation of the Cyclic Peptide Database (Cyclome930)

To enable large-scale analysis of cyclic peptides, we constructed a comprehensive and curated database by aggregating experimentally determined cyclic peptide structures from multiple public resources, including the Protein Data Bank (PDB),^32^ ConoServer,^31^ CyBase,^27^ and Cyclicpepedia.^28^ Structures deposited up to 2025 were considered. An initial collection of 1,059 cyclic peptide–containing structures was assembled, comprising entries determined by X-ray crystallography, nuclear magnetic resonance (NMR), and cryo-electron microscopy. Because many deposited structures contain multiple peptide chains, each structure was decomposed into its constituent peptide chains, yielding a total of 1,734 individual chains. Chains were filtered to retain only cyclic peptides based on explicit cyclization annotations or the presence of covalent linkages consistent with end-to-end (e2e) or side-chain–mediated cyclization (s2e, s2s, e2e+s2s, s2e+s2s). Redundant entries arising from identical sequences appearing in multiple structures were consolidated, resulting in a final non-redundant dataset of 930 unique cyclic peptides, referred to as Cyclome930.

Sequence lengths of peptides in Cyclome930 were analyzed to assess dataset diversity. The database spans a broad range of peptide lengths, with the majority of entries falling below 40 residues, while still including longer cyclic peptides approaching 200 residues. This distribution highlights the structural and functional diversity represented in the curated dataset and supports its suitability for downstream statistical and machine-learning analyses. Each cyclic peptide entry in Cyclome930 was annotated with detailed metadata extracted from primary structure records and associated literature. Annotations include : source organism, covering a wide range of natural producers, functional classification, where available, method of structure determination (e.g., X-ray crystallography, NMR), peptide origin, distinguishing wild-type, mutant, and synthetic constructs etc. These annotations enable stratified analyses of cyclic peptides across biological origin, experimental methodology, and functional classes.

### Cyclicity-aware -sequence Similarity Calculator

Accurate assessment of sequence similarity among cyclic peptides requires explicit consideration of cyclization topology, as conventional linear sequence alignment approaches assume fixed N- and C-termini and therefore underestimate similarity for circular peptides. To address this limitation, we developed a cyclization-aware sequence similarity calculation framework that adapts the alignment strategy based on the type of cyclization present in the peptide.

#### End-to-End (e2e) Cyclized Peptides

For peptides cyclized through an end-to-end (e2e) linkage, rotational symmetry renders the linear sequence representation ambiguous. To account for this, the template sequence is duplicated and concatenated in the N→C direction to generate a true template (*T-template*) that spans two full sequence lengths. Sequence similarity between a query peptide and the T-template is then evaluated using local alignment, allowing the query sequence to align optimally across all possible cyclic rotations (**Figure 4**).

#### Side-to-End (s2e) and Side-to-Side (s2s) Cyclized Peptides

For peptides cyclized through side-to-end (s2e) or side-to-side (s2s) linkages, the linear sequence order remains well-defined and adequately represents the underlying peptide topology. In these cases, the original template sequence is used directly for similarity calculation without duplication or resegmentation

#### End-to-End with Side-to-Side (e2e+s2s) Cyclized Peptides

For peptides containing both end-to-end cyclization and one or more side-to-side bridges (e2e+s2s), multiple candidate T-templates are constructed to capture the increased topological complexity. First, the full sequence is duplicated and concatenated in the N→C direction, as described for e2e peptides. In addition, for each s2s bridge, the sequence is segmented from the N-terminus to the first residue involved in the bridge, and this fragment is concatenated with the segment beginning at the second bridging residue. This procedure generates alternative T-templates corresponding to different cyclic traversal paths. Sequence similarity between the query peptide and each T-template is computed using local alignment, and the maximum similarity score across all T-templates is reported.

#### Side-to-End with Side-to-Side (s2e+s2s) Cyclized Peptides

For peptides containing both side-to-end and side-to-side cyclization (s2e+s2s), T-template construction follows the same segmentation and recombination strategy described for e2e+s2s peptides. Similarity scores are computed between the query sequence and each constructed T-template, and the highest score is retained as the final similarity measure.

#### Similarity Scoring

Across all cyclization classes, similarity is evaluated using local sequence alignment to accommodate partial matches and motif-level conservation. By explicitly accounting for cyclic topology and multiple possible traversal paths, this approach ensures that similarity scores reflect true sequence relatedness rather than artifacts of linearization.

### Cyclization classification

The cyclization topology of each peptide in the dataset was systematically determined using a geometry-based classification scheme derived from inter-residue distance calculations. Specifically, distances between relevant backbone and side-chain atoms were computed from experimentally determined structures to identify covalent linkages consistent with different cyclization modes. Classification criteria and distance thresholds are summarized in **Table 3**.

**Table 3.**
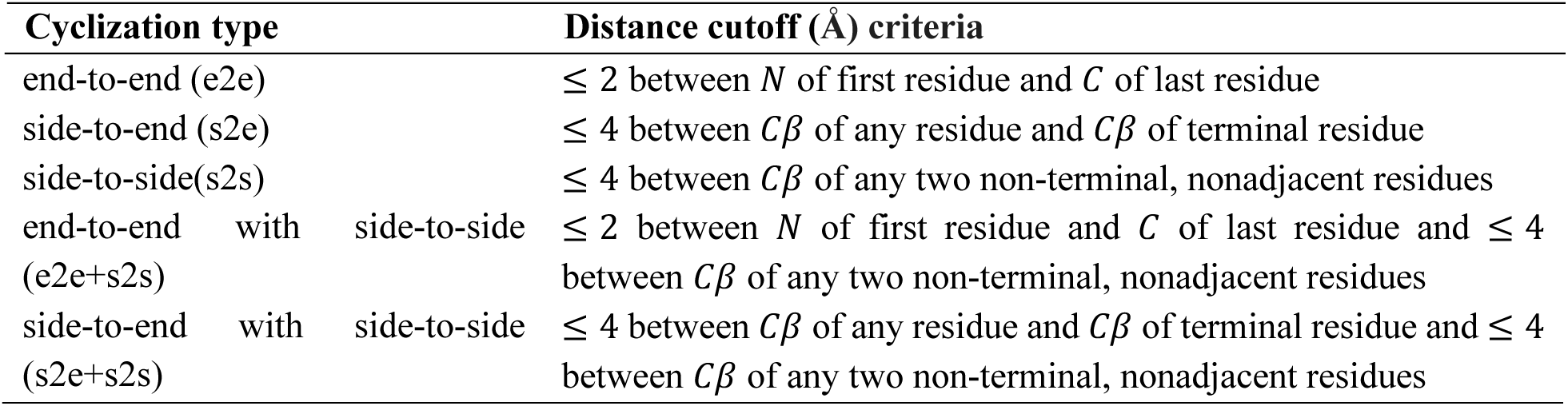
Cyclization types and their corresponding distance cutoff criteria.

| Cyclization type | Distance cutoff (Å) criteria |
| --- | --- |
| end-to-end (e2e) | $\leq 2$ between $N$ of first residue and $C$ of last residue |
| side-to-end (s2e) | $\leq 4$ between $C\beta$ of any residue and $C\beta$ of terminal residue |
| side-to-side(s2s) | $\leq 4$ between $C\beta$ of any two non-terminal, nonadjacent residues |
| end-to-end with side-to-side (e2e+s2s) | $\leq 2$ between $N$ of first residue and $C$ of last residue and $\leq 4$ between $C\beta$ of any two non-terminal, nonadjacent residues |
| side-to-end with side-to-side (s2e+s2s) | $\leq 4$ between $C\beta$ of any residue and $C\beta$ of terminal residue and $\leq 4$ between $C\beta$ of any two non-terminal, nonadjacent residues |

This distance-based classification framework enables consistent and automated assignment of cyclization types across a large and structurally diverse set of cyclic peptides, providing a foundation for downstream sequence analysis, structural comparison, and machine-learning model development.

### Temperature ramped molecular dynamics simulations to generate melting point priors of cyclic peptides

As experimental melting points for cyclic peptides are very scarce, we resorted to a computational method to generate melting point indicators from structural changes in these peptides. We performed molecular dynamics simulations for all cyclic peptides in Cyclome930 (see **Supplementary Data 1** for the list) using Gromacs version 2023.2 molecular simulation software.^33,34^ Our simulation framework solvates the cyclic peptide in a cubic water box, followed by energy minimization employing the steepest descent method.^35^ This was followed by a 100-ps equilibration phase in the canonical ensemble at 298 K, with restraint force applied solely to the heavy atoms of the protein structure. The NVT stabilization phase was succeeded by an isobaric-isothermal (NPT) equilibration, rigorously maintained at a pressure of 1 bar and a temperature of 298 K, employing the Parrinello-Rahman barostat.^36^ The subsequent production MD trajectories spanned 10 ns under the aforementioned thermodynamic conditions, integrating at a timestep of 2 fs. Both the equilibration and production phases utilized the inherent leap-frog integrator embedded within the GROMACS suite.

For the temperature- replica exchange molecular dynamics (T-REMD) simulations, the last frame of the single temperature production run is used as the first frame of simulations. The number of replicas for each cyclic peptide simulation box was calculated using *Virtual Chemistry – remd-temperature-generator*,^37^ with an exchange probability of ∼ 0.2 between replicas. A temperature range of 298 - 400 K is set for T-REMD simulations. The dynamics of each cyclic peptide is performed for 100 ns for each replica.

After the completion of the simulations, the frames in replicas are sorted based on the average temperature/energy, before it was further analyzed for priors of melting-like transitions. Trajectory analyses such as ramachandran angles (Δ*ϕ*, Δ*ψ*), root mean square fluctuation (RMSF) of peptide backbone, radius of gyration (g_r_) of the peptide backbone along all the three axes are done to understand the change in peptide structure on heating. The temperature of the replica in which a drastic change i.e., 20 % increase or decrease in one of these parameters with respect to the initial structure or 10 % change in two or more of these parameters with respect to the initial structure is considered its melting point (STop2Melt). Another criterion for STop2Melt is that the change in structure does not revert back to its original structure in any of the replicas that has higher temperature than that of the melting point.

### STop2Melt predictor model

#### Input data and target variable

We trained a supervised regression model to predict the REMD-derived melting-point proxy (STop2Melt) for Cyclome930. Each entry contains (i) peptide sequence, (ii) cyclization class, (iii) cyclization site, (iv) STop2Melt labels. The overall framework for STop2Melt predictor is given in **Figure 9a**.

#### Sequence representation using ESMc embeddings

To featurize cyclic peptides, we used pretrained ESMc protein language model embeddings (300M; EvolutionaryScale/esmc-300m-2024-12).^38^ For each peptide sequence of length *L*, ESMc produced a per-residue embedding matrix. We converted variable-length embeddings into fixed-length sequence descriptors by applying both mean and max pooling across the residue dimension and concatenated them to form the primary sequence feature vector. This pooling strategy preserves global sequence information while enabling standard regression models to operate on fixed-size inputs (**Figure 9a**).

#### Encoding cyclic topology (“cyclic offset” summary features)

To incorporate cyclicity awareness of sequence, we derived a topology descriptor from the peptide’s cyclization pattern. For example, if a sequence is cyclized at `GLY1-ASN30` and `CYS4-CYS20` we parsed the residue indices involved in each covalent linkage here (1, 30) and (4,20). We then constructed a residue graph with (i) backbone edges (*i*, *i* + 1) and (ii) additional edges defined by the cyclization links. From this graph D, we computed a neighbor matrix of all residues. From D we extracted two families of cyclic-topology summary features: <u>(1) Distance histogram:</u> the normalized frequency of each discrete distance bucket 0. . *d_max_* across all residue pairs, providing a compact description of global topological compactness, <u>(2) Graph degree statistics:</u> residue degrees were computed as deg(*i*) = ∑*_j_* 1[*D_ij_* = 1], and we included {mean,std,min,max} of deg(*i*) as global descriptors of connectivity.

#### Cyclization-class (topology) conditioning

Cyclization classes (e2e, s2s, e2s, e2e+s2s, e2s+s2s) were incorporated as an additional categorical input. We one-hot encoded the cyclization class label and appended it to the feature vector to allow the regressor to learn class-dependent relationships between sequence/topology and STop2Melt. These features were concatenated with the pooled ESM representation:

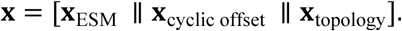

#### Cluster-aware train/validation/test partitioning

To prevent overly optimistic performance due to near-duplicate sequences and shared cyclization patterns across splits, we used cluster-aware data partitioning. Each peptide was assigned to a “cluster” defined by the tuple:

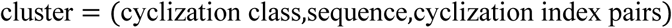

We then performed a cluster-level split into **train/validation/test** sets (default 70/15/15 by cluster count), ensuring no cluster appeared in more than one partition. We enforced that all cyclization classes were represented in each partition by allocating at least one cluster per class to the validation and test sets prior to filling the remaining clusters to meet the desired split fractions.

#### Regression models and training protocol

We evaluated multiple regression algorithms using the same features and identical cluster-aware splits: Ridge, Lasso, ElasticNet, SVR (RBF kernel), K-Nearest Neighbors, Random Forest, ExtraTrees, Gradient Boosting, and a feed-forward MLP regressor. For models sensitive to feature scaling (Ridge/Lasso/ElasticNet/SVR/KNN/MLP), features were standardized (zero mean, unit variance) using parameters fit on the training set only. Tree-based models were trained directly on the raw concatenated features.

#### Model evaluation

Performance was reported on the held-out test set using standard regression metrics: mean absolute error (MAE), root-mean-square error (RMSE), and coefficient of determination (R²). Hyperparameters were kept consistent across splits and models as implemented in our benchmarking pipeline; model selection can be performed based on validation MAE, with final reporting on the unseen test set to estimate generalization under cluster-aware conditions.

#### CritiCL- multiclass classification model for critical mineral screening

The CritiCL classifier was developed as a multiclass prediction framework to infer the metal-binding preference of protein/peptide sequences. Four supervised machine learning algorithms were trained on linear peptides and proteins that bind to Ln^3+^, Co^2+^, Ni^2+^, Mn^2+^ or others (any other transition metal). The overall framework for CritiCL is given in **Figure 10**. The classification frameworks evaluated are: Random Forest (RF), Extreme Gradient Boosting (XGBoost), Support Vector Machine (SVM), and Multilayer Perceptron (MLP). These models were selected to represent complementary learning paradigms, including ensemble decision trees, kernel-based methods, and neural networks. All models were trained to predict one of five metal-binding classes: Ln^3+^, Co^2+^, Ni^2+^, Mn^2+^or others Class labels were encoded using a categorical label encoder, and the models were trained using sequence embedding vectors that are compatible to cyclicity aware embeddings as input features (similar to STop2Melt predictor). Cyclome930 was tested on CritiCL (zero shot) to predict which sequences can potentially bind to critical minerals such as Ln^3+^and Co^2+^.

#### Evaluation Metrics

Classifier performance was assessed using multiple complementary metrics to capture different aspects of predictive performance. Overall accuracy and macro-averaged F1 score were used to evaluate classification performance across all classes. The Matthews correlation coefficient (MCC) was calculated to provide a balanced measure of prediction quality for multiclass classification problems. Additionally, receiver operating characteristic area under the curve (ROC-AUC) values were computed using a one-versus-rest approach to assess class separability.

#### Screening of Cyclic Peptide Candidates

After training on linear peptides/proteins, the CritiCL classifier was applied to Cyclome930. For each sequence, the model generated probability scores corresponding to Ln^3+^, Co^2+^, Ni^2+^, Mn^2+^ or others binding classes. These scores were used to rank peptides according to their predicted affinity for critical mineral ions, enabling rapid in silico screening of peptide libraries for candidate metal-binding sequences.

## Contributions

Conceptualization: R.C., K.A.S.

Methodology: K.A.S., H.G, J.Y, R.C

Investigation: K.A.S., H.G

Visualization: K.AS, R.C, C.P.H.T, R.D

Funding acquisition: R.C.

Project administration: R.C.

Supervision: R.C.

Writing: original draft: K.A.S, R.C

Writing: review & editing: K.A.S., H.G., V.S.R., R.C.

Webserver: C.P.H.T, R.D

## Data and Code availability

The Cyclome930 dataset and cyclicity aware sequence similarity test inference is available for download at https://cyclome930.studio/, inference code for STop2Melt is available at Stop2Melt_inference and the inference code for CritiCL is available at CritiCL_inference.

## Acknowledgements

This research is partially supported by the U.S. Department of Energy, Office of Science, Office of Biological and Environmental Research, Biological Systems Science Division, 2025 Lab Critical Materials and Minerals (CMM) Pilot Program to R.C. Ames National Laboratory is operated for the U.S. Department of Energy by Iowa State University under Contract No. DEAC02-07CH11358. Cyclome930 data curation was supported by the Critical Materials Innovation Hub, an Energy Innovation Hub funded by the US Department of Energy, Office of Critical Minerals and Energy Innovation (CMEI), Advanced Materials and Manufacturing Technologies Office (AMMTO) - under contract DEAC52–07NA27344 (LLNL-JRNL-857236). CritiCL data collection was also supported by Ames Laboratory LDRD to R.C.

