## Supplemental Material for "Cyclome: Large-scale replica-exchange dynamics of 930 cyclic peptide reveal thermal stability and critical metal-binding behavior"

### Supplementary Figures

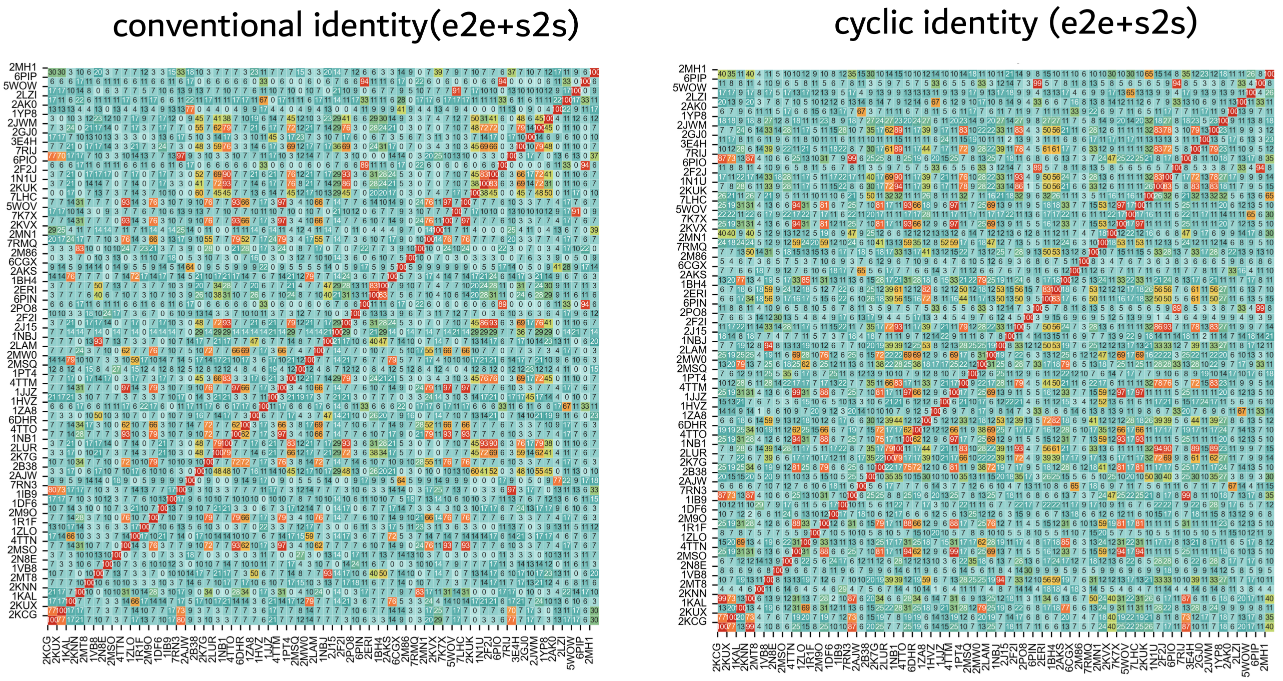


**S. Figure 1.** Pairwise identity heatmaps illustrating sequence/structure relationships across the e2e+s2s cyclic peptide dataset using conventional scheme and cyclicity aware scheme. Each matrix encodes all-against-all similarities, with warmer colors indicating higher similarity and cooler colors indicating lower similarity. The strong diagonal corresponds to self-similarity, while the off-diagonal patterns reflect inter-peptide relationships. Compared to the baseline representation (left), the enhanced representation (right) exhibits improved contrast and organization in identity patterns, suggesting better discrimination of structurally and topologically related cyclic peptides.

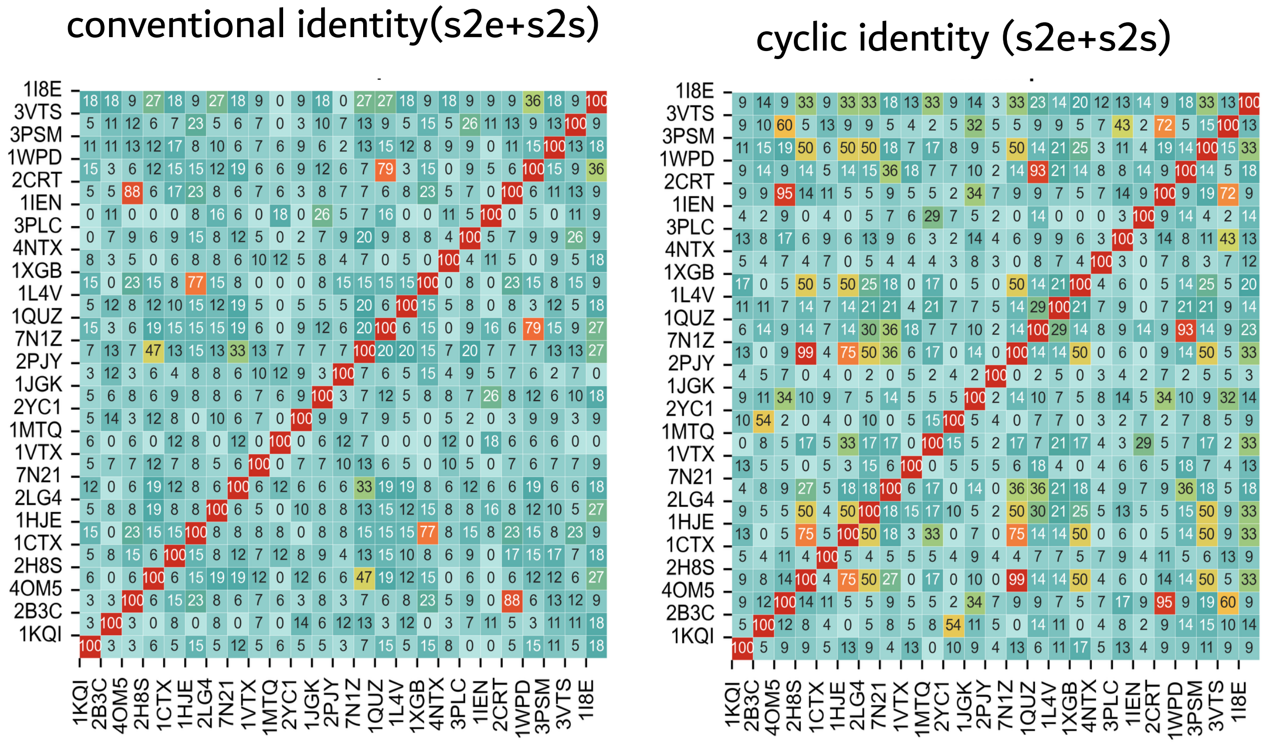


**S. Figure 2.** Pairwise identity matrices of s2e+s2s cyclic peptides highlighting sequence/structure relationships across the dataset under two alignment strategies. Each heatmap represents all-against-all similarity scores, with warmer colors indicating higher similarity and cooler colors indicating lower similarity. The diagonal reflects self-identity, while off-diagonal regions capture inter-peptide relationships. Compared to the conventional alignment (left), the cyclicity-aware alignment (right) exhibits sharper contrast and enhanced detection of related peptide clusters, demonstrating improved sensitivity to cyclic topology and sequence equivalence.

a

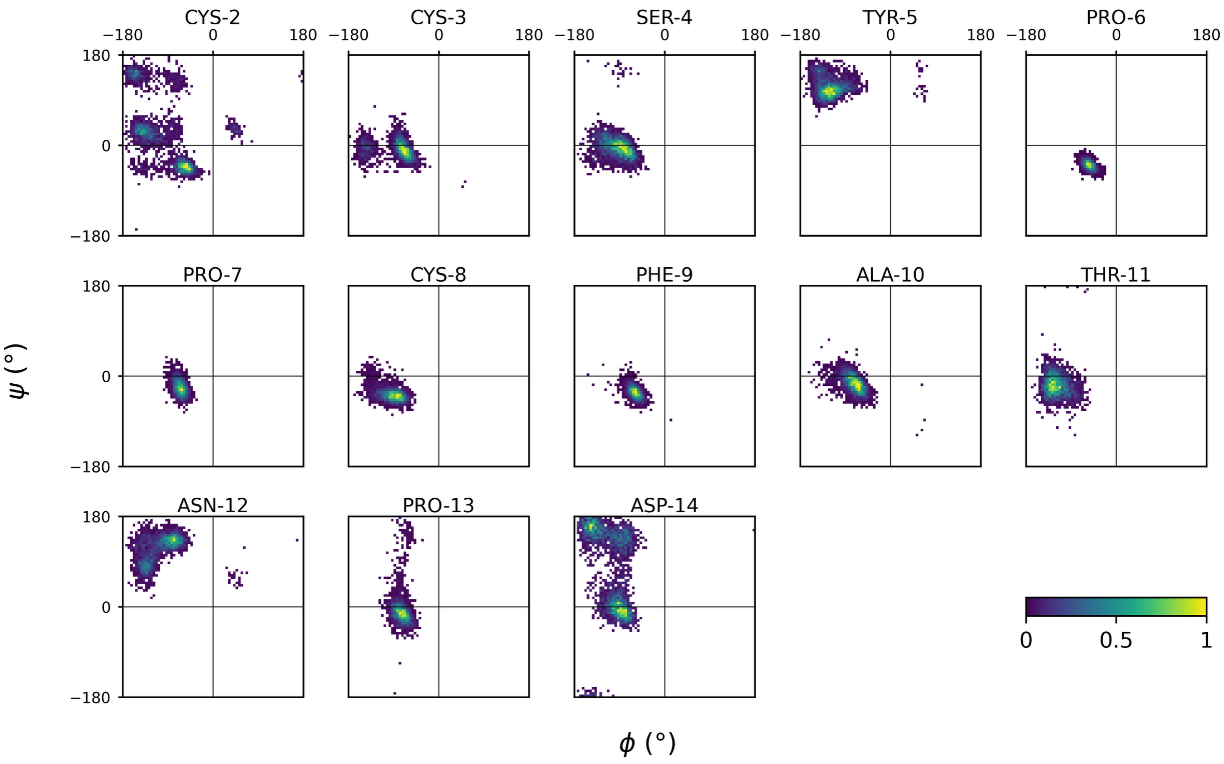


b

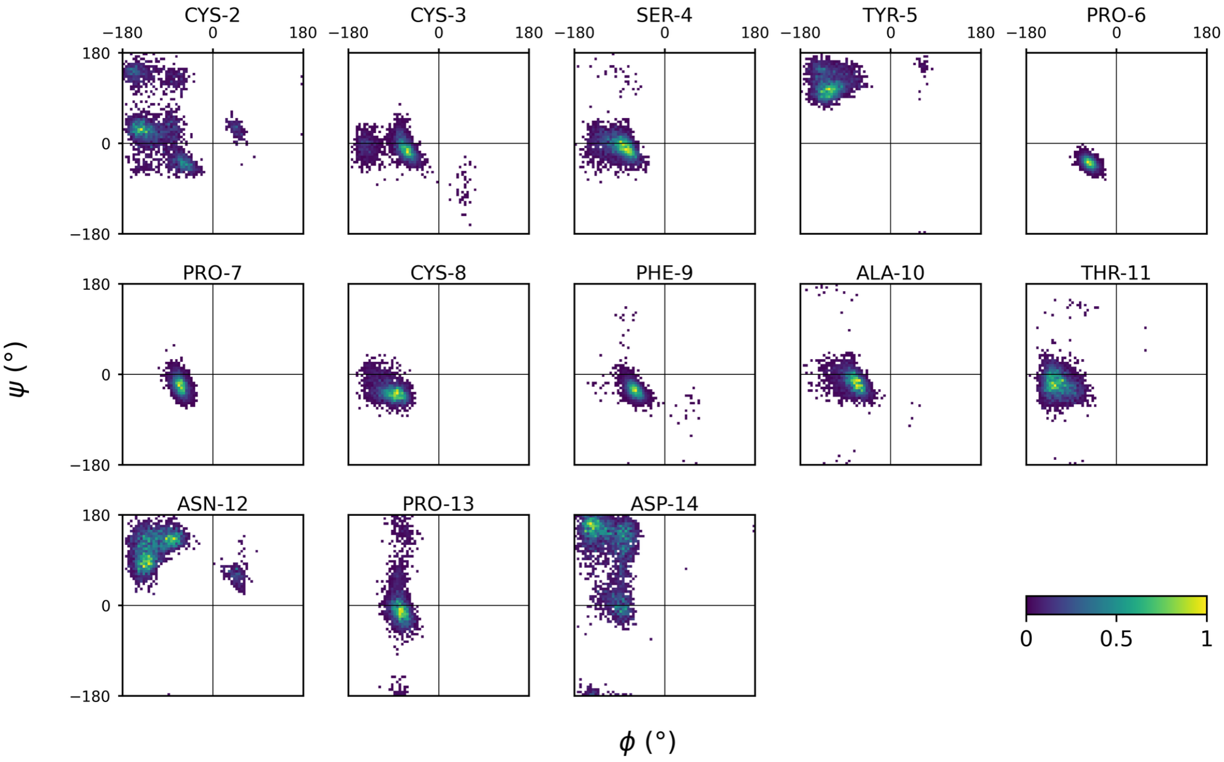


**S. Figure 3.** Residue-wise Ramachandran density distributions (ϕ, ψ) illustrating conformational sampling across the residues under two thermodynamic regimes. Each panel corresponds to an individual residue, with color intensity indicating normalized population density. (**a**) represents conformational ensembles below the STop2Melt transition, characterized by more localized and well-defined dihedral populations, reflecting structurally constrained states. In contrast, (**b**) corresponds to ensembles at STop2Melt, where broader and more diffuse distributions are observed, indicating increased conformational flexibility and sampling of higher-energy states. Notably, residue-specific variations highlight differential flexibility across the peptide.

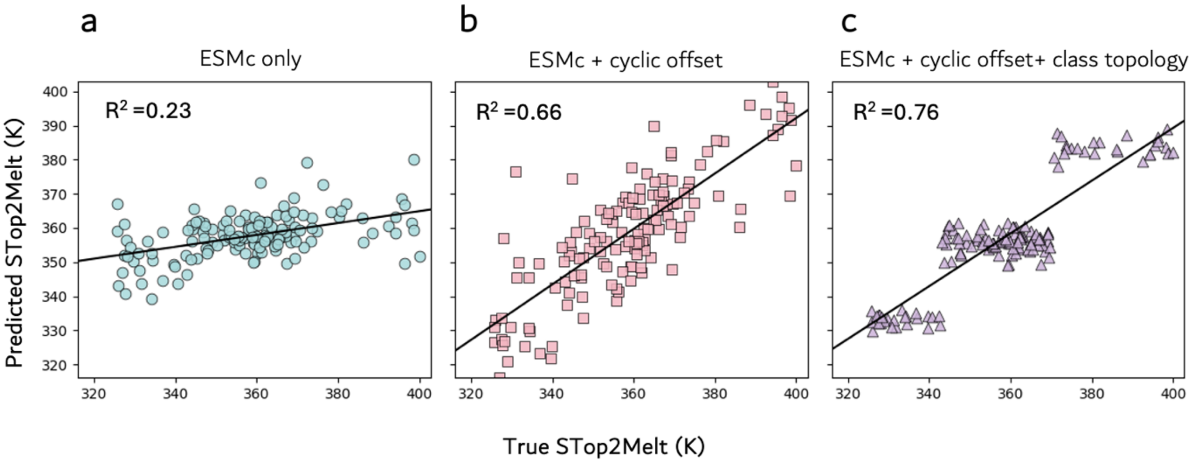


**S. Figure 4.** Comparison of true vs predicted values for STop2Pred cyclic peptides across progressively enriched feature representations. (**a**) Model trained using ESMc embeddings alone shows weak correlation with experimental values ($R^2$ = 0.23), indicating limited capture of thermodynamic determinants. (**b**) Incorporation of cyclic offset features substantially improves predictive performance ($R^2$ = 0.66), highlighting the importance of encoding cyclic topology. (**c**) Further inclusion of cyclization class topology descriptors yields the highest agreement with true values ($R^2$ = 0.76), demonstrating that explicit structural topology information enhances model accuracy. Solid black lines represent linear regression fits for each model.

**S. Table 1.** Performance metrics for STop2Melt model with ESMc only embedding

| Model | Val_MAE | Val_RMSE | Val_$R^2$ | Test_MAE | Test_RMSE | Test_$R^2$ |
|---|---|---|---|---|---|---|
| **SVR-RBF** | 11.27 | 14.68 | 0.22 | 11.39 | 15.28 | 0.23 |
| **GradientBoosting** | 11.90 | 15.43 | 0.14 | 11.63 | 15.45 | 0.21 |
| **RandomForest** | 11.30 | 14.33 | 0.25 | 11.69 | 15.33 | 0.22 |
| **ExtraTrees** | 10.96 | 14.05 | 0.28 | 11.77 | 15.46 | 0.21 |
| **KNN** | 11.48 | 14.94 | 0.19 | 12.30 | 16.17 | 0.13 |
| **Ridge** | 16.50 | 22.0 | -0.76 | 18.27 | 24.80 | -1.04 |
| **ElasticNet** | 17.33 | 22.62 | -0.86 | 18.78 | 25.11 | -1.09 |
| **Lasso** | 19.42 | 25.64 | -1.39 | 20.24 | 27.02 | -1.42 |
| **MLP** | 23.35 | 33.97 | -3.19 | 22.70 | 31.50 | -2.29 |

**S. Table 2.** Performance metrics for STop2Melt model with ESMc+ cyclic offset embedding

| Model | Val_MAE | Val_RMSE | Val_$R^2$ | Test_MAE | Test_RMSE | Test_$R^2$ |
|---|---|---|---|---|---|---|
| **ExtraTrees** | 7.417 | 82.46 | 0.70 | 8.43 | 102.72 | 0.66 |
| **RandomForest** | 7.51 | 84.24 | 0.69 | 8.64 | 112.44 | 0.63 |
| **GradientBoosting** | 7.69 | 89.42 | 0.68 | 8.86 | 129.27 | 0.57 |
| **SVR-RBF** | 10.93 | 198.51 | 0.28 | 10.94 | 212.48 | 0.30 |
| **KNN** | 11.40 | 219.49 | 0.20 | 12.01 | 249.90 | 0.17 |

| | | | | | | |
|---|---|---|---|---|---|---|
| **Ridge** | 13.24 | 336.21 | -0.22 | 13.58 | 325.92 | -0.08 |
| **ElasticNet** | 14.03 | 389.51 | -0.41 | 14.48 | 369.11 | -0.22 |
| **Lasso** | 16.58 | 515.10 | -0.87 | 17.43 | 520.98 | -0.73 |
| **MLP** | 21.65 | 1057.24 | -2.84 | 20.59 | 834.62 | -1.77 |

**S. Table 3.** Performance metrics for STop2Melt model with ESMc+ cyclic offset +topology embedding

| Model | Val_MAE | Val_RMSE | Val_$R^2$ | Test_MAE | Test_RMSE | Test_$R^2$ |
|---|---|---|---|---|---|---|
| **RandomForest** | 6.67 | 63.99 | 0.77 | 7.20 | 71.14 | 0.76 |
| **GradientBoosting** | 6.91 | 71.18 | 0.74 | 7.29 | 75.20 | 0.75 |
| **ExtraTrees** | 6.94 | 71.91 | 0.74 | 7.63 | 82.16 | 0.73 |
| **SVR-RBF** | 10.75 | 190.37 | 0.31 | 10.68 | 200.57 | 0.34 |
| **KNN** | 11.36 | 214.12 | 0.22 | 11.98 | 247.80 | 0.18 |
| **Ridge** | 12.76 | 304.60 | -0.11 | 13.38 | 299.95 | 0.01 |
| **ElasticNet** | 13.73 | 357.33 | -0.30 | 14.52 | 350.02 | -0.16 |
| **Lasso** | 14.76 | 416.56 | -0.51 | 17.23 | 521.12 | -0.73 |
| **MLP** | 22.59 | 1237.32 | -3.49 | 20.10 | 854.29 | -1.83 |
